# Molecular Logic of Spinocerebellar Tract Neuron Diversity and Connectivity

**DOI:** 10.1101/516435

**Authors:** Myungin Baek, Vilas Menon, Thomas M. Jessell, Adam W. Hantman, Jeremy S. Dasen

## Abstract

Coordinated motor behaviors depend on feedback communication between peripheral sensory systems and central circuits in the brain and spinal cord. Relay of muscle and tendon-derived sensory information to the CNS is facilitated by functionally and anatomically diverse groups of spinocerebellar tract neurons (SCTNs), but the molecular logic by which SCTN diversity and connectivity is achieved is poorly understood. We used single cell RNA sequencing and genetic manipulations to define the mechanisms governing the molecular profile and organization of SCTN subtypes. We found that SCTNs relaying proprioceptive sensory information from limb and axial muscles are generated through segmentally-restricted actions of specific *Hox* genes. Loss of *Hox* function disrupts SCTN subtype-specific transcriptional programs, leading to defects in the connections between proprioceptive sensory neurons, SCTNs, and the cerebellum. These results indicate that Hox-dependent genetic programs play essential roles in the assembly of the neural circuits required for proprioception.

## Introduction

Relay of muscle-derived sensory information from the periphery to the CNS is essential for coordinating motor output during behavior, and plays essential roles during motor learning and adaptation^1,2^. The role of proprioception in motor control has been investigated in animal studies where sensory neurons have been genetically or surgically ablated, as well as in sensory neuropathies that disrupt proprioceptive feedback^3^. While basic motor functions such as walking and reaching are retained, loss of proprioception causes severe defects in limb coordination. In humans with sensory deficits, the ability to move the arm is maintained, but characterized by the inability to predict and correct errors^4,5^. Ablation of hindlimb proprioceptive input leads to a loss of inter-joint limb coordination, as well as defects in the ability of animals to adapt locomotor behaviors when confronted with uneven terrains^6–8^.

Muscle and joint-derived sensory information is relayed to the CNS through specialized classes of proprioceptive sensory neurons (pSNs) that connect peripherally with muscle spindles and Golgi tendon organs^9^. Centrally pSNs establish connections with diverse arrays of neuronal subtypes including spinal motor neuron (MNs), local circuit interneurons, and ascending projection neurons. Ascending pathways relay information related to muscle contractile status to higher brain centers, including the cerebellum. Proprioceptive sensory streams are transmitted to the cerebellum through neurons that project along the spinocerebellar and cuneocerebellar tracts^1,10^. Spinal projections originating from spinocerebellar tract neurons (SCTNs) terminate as mossy fibers and constitute a major source of input to cerebellar granule cells.

Anatomical tracing studies in mammals indicate that SCTNs comprise up to a dozen distinct subtypes which are located at discrete positions along the rostrocaudal axis of the spinal cord^11–14^. Electrophysiological studies, predominantly in cats and rats, have shown that each SCTN type is targeted by pSNs that innervate specific muscle groups. For example, neurons within Clarke’s column relay proprioceptive information from hindlimb muscles, the central cervical nucleus from the neck, and Stilling’s sacral nucleus from the tail^10,15,16^. In contrast to the more selective central connections between pSN central afferents and MN pools, neurons within Clarke’s column appears to receive sensory inputs from multiple, and often functionally antagonistic, muscle types^17,18^. This has led to the proposal that the information relayed from pSNs to Clarke’s column neurons provides more global information about limb parameters, such as direction of limb movement and orientation, as opposed to muscle-specific features^10^. In addition to input from pSNs, neurons within Clarke’s columns receive direct excitatory and indirect inhibitory input from corticospinal neurons^19^. The coincidence of cortical and muscle-derived inputs suggests that SCTNs function as local hubs that integrate and process sensory and motor information.

Despite progress in elucidating the anatomical organization and physiological features of SCTNs, the molecular basis for their subtype diversification and connectivity is largely unknown. In principle, SCTN diversification could employ the same developmental mechanisms that have been defined for other neuronal classes, such as spinal MNs. All spinal MNs arise from a single progenitor domain, but give rise to dozens of topographically organized muscle-specific subtypes^20^. This diversity is established through the activities of Hox transcription factors along the rostrocaudal axis. *Hox* genes are expressed by multiple neuronal populations within the hindbrain and spinal cord, suggesting a broader role neuronal specification. Although recent studies have implicated *Hox* function during the differentiation of interneurons in the ventral spinal cord^21,22^, the identity of their downstream target effectors and potential roles in sensory-motor circuit assembly have not been investigated.

We used single cell RNA sequencing to define the molecular signatures of SCTNs generated at cervical and thoracic levels of the spinal cord. We show that the specification of SCTNs relies on segmental-level specific activities of Hox transcription factors, and loss of *Hox* gene function transforms the molecular profiles and connectivity patterns of SCTN subtypes. These results indicate that the specification of SCTNs relies on the same developmental programs used to generate spinal MN subtypes, suggesting a common transcriptional strategy drives cell type diversification across multiple neuronal classes.

## Results

### Organization and input specificity of SCTNs

To dissect the molecular profiles of SCTN subtypes, we first used retrograde tracing from the cerebellum to map the position of SCTNs along the rostrocaudal axis of the spinal cord. We injected Alexa555 conjugated cholera toxin B (CTB) into the cerebellum of P4 mice and allowed SCTNs to be labeled for 2 days. Whole-mount staining of the spinal cord labeled specific subsets of neurons along the rostrocaudal axis (Fig. 1a). Prominent columns of neurons were found near the midline of rostral cervical, thoracic and rostral lumbar levels; and more laterally-positioned columns at caudal lumbar and sacral levels. More scattered SCTN populations were found throughout the entire length of spinal cord. We mapped the distribution of SCTNs within specific spinal segments and generated contour maps of SCTN densities at cervical, thoracic, lumbar, and sacral levels (Fig. 1b,c). Consistent with previous studies, four prominent clusters of SCTNs were labeled, including the central cervical nucleus (CCN) at rostral cervical levels, Clarke’s column (CC) neurons extending from thoracic to rostral lumbar levels, spinal border cells (SBC) at lumbar levels, and Stilling’s nucleus (SSN) at sacral levels^13–15^. We also identified SCTNs showing more distributed patterns at cervical levels in Rexed lamina (L)V, LVI, and LVII, and at lumbar levels in LV, LVII, and LVIII (Fig. 1c). Collectively, these tracing data identify 10 major groups of SCTNs in early postnatal mice (Fig. 1e, and Supplementary Fig. 1).

**Fig. 1.**
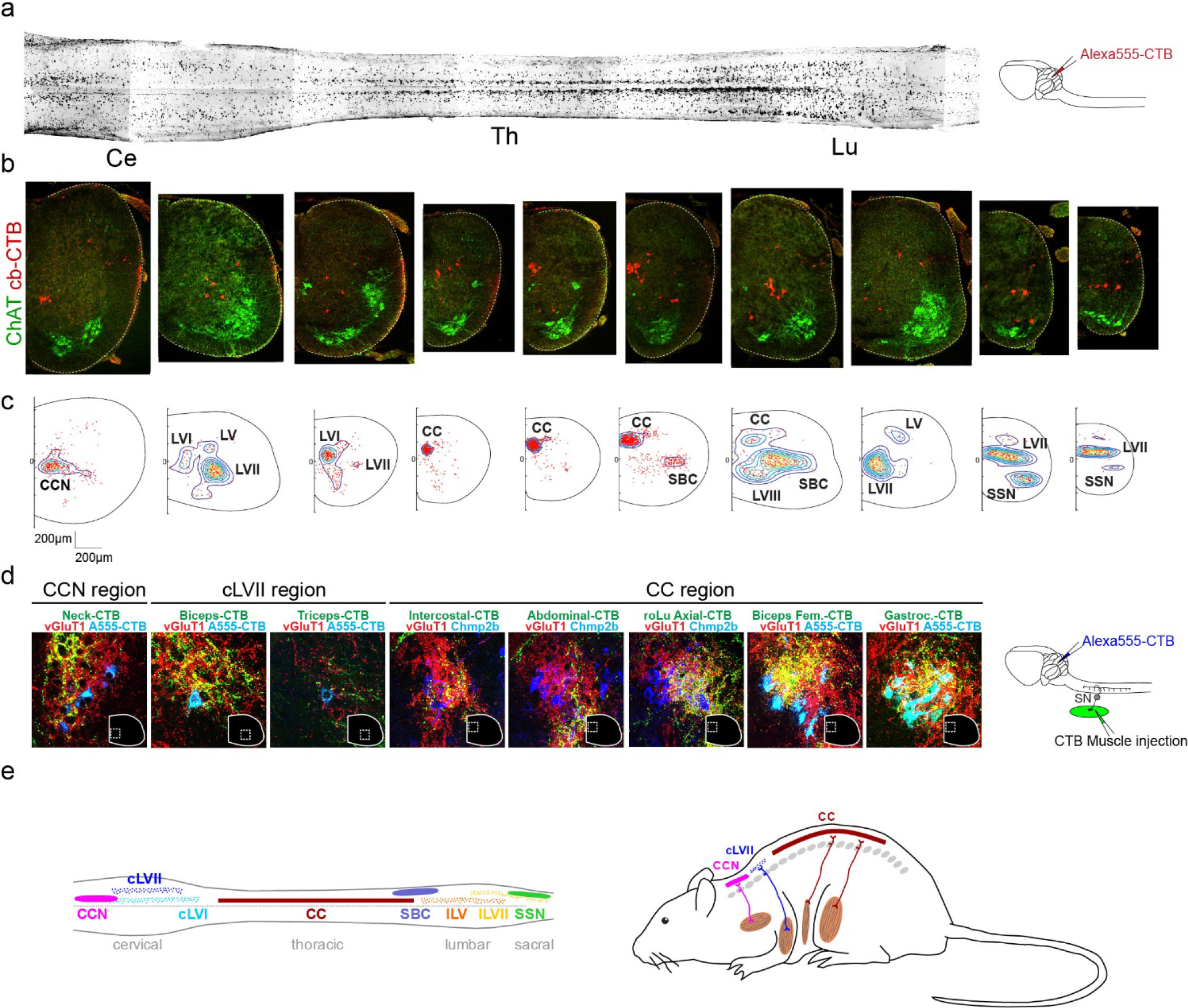
Distribution and muscle-specific inputs of SCTNs. (a-c) SCTNs were retrogradely labeled by injection of Alexa-555 conjugated cholera toxin B subunit (Alexa555-CTB) into the cerebellum at postnatal day 4 (P4) and analyzed at P6. (a) Whole mount of Alexa555-CTB labeled P6 mouse spinal cord. Injection schematic is shown on the right. Ce, cervical; Th, thoracic; Lu, lumbar (b) Top: Alexa555-CTB labeled spinal cord sections. Shown are the matched regions to the whole mount spinal cord. Last two sections are from sacral regions. Cb-CTB, Alexa555-CTB signal from the cerebellum injection. (c) Density plots of labeled SCTNs. Contour plots were generated from spinal cords of n=6 mice. Number of cells in each section, from left to right (rostral cervical to sacral) is 299, 165, 251, 241, 376, 662, 266, 78, 161, and 92 cells. Distance, μm. red dots, labeled individual SCTNs. (d) Sensory inputs to SCTNs were traced by CTB injection into indicated target muscle. Shown are the magnified images of regions demarked by white dashed lines. VGluT1 labels pSN terminals; A555-CTB labels traced SCTNs, Chmp2b marks Clarke’s column neurons (found in Allen brain atlas database). Injection schematic is shown on the right. (e) Summary of SCTN organization in mouse.

SCTNs are essential for relaying proprioceptive sensory information from muscle to cerebellum, but the muscle-specific inputs that SCTNs receive are largely unmapped in mouse. We examined the source of inputs from proprioceptive sensory neurons to SCTNs by injection of CTB into specific muscles, while in parallel labeling SCTNs with either cerebellar retrograde tracing or using SCTN-restricted molecular markers. Selectivity of proprioceptive inputs was further delineated by localization with VGluT1, which labels the presynaptic boutons of pSNs^23,24^. This analysis revealed that SCTNs receive input from discrete muscle types, and are consistent with studies in rat and cat^10,15,24,25^. Rostral cervical CCN neurons receive inputs from pSNs innervating neck muscles, caudal cervical LVII SCTNs receive input from forelimb muscle, while thoracic/upper lumbar CC neurons receive input from axial and hindlimb muscles (Fig. 1d). Inputs to SBC neurons were not labeled through any of the muscle injections we attempted, and did not contain VGluT1+ presynaptic boutons, as previously reported (data not shown)^24^. These results indicate that specific populations of SCTNs can be delineated by their rostrocaudal position, settling location, and the source of their inputs from specific muscle groups.

### Molecular profiling of SCTNs at cervical and thoracic levels

To determine whether SCTN subtypes can be distinguished by differences in molecular profiles, we performed RNA sequencing (RNAseq) on retrogradely labelled and individually isolated SCTNs from cervical and thoracic levels (Fig. 2a). To obtain high sequencing depth we first performed RNAseq on pools of labeled SCTNs. We collected 4 pools, each containing ~200 cervical SCTNs, and 4 pools of ~350 thoracic SCTNs. We identified 1768 genes that were enriched in cervical SCTNs and 495 genes enriched in thoracic SCTNs (>2-fold change; FDR<0.05) (Fig. 2b). Differentially expressed genes included effector molecules with implications for neural function including ion channels, neuropeptide receptors, and neurotransmitter transporters (Fig. 2c). For example, selective expression of neuropeptides and associated proteins was found in cervical SCTNs (e.g. *NPY*, *Tac1*,*Pnoc*, *pdyn*, *qrfp*,*scg2*) and thoracic SCTNs (*NTS*), suggesting that SCTN subtypes differentially release more than one neuromodulator. This dataset will be useful for testing hypotheses about anatomical and physiological differences between cervical and thoracic SCTN populations.

**Fig. 2.**
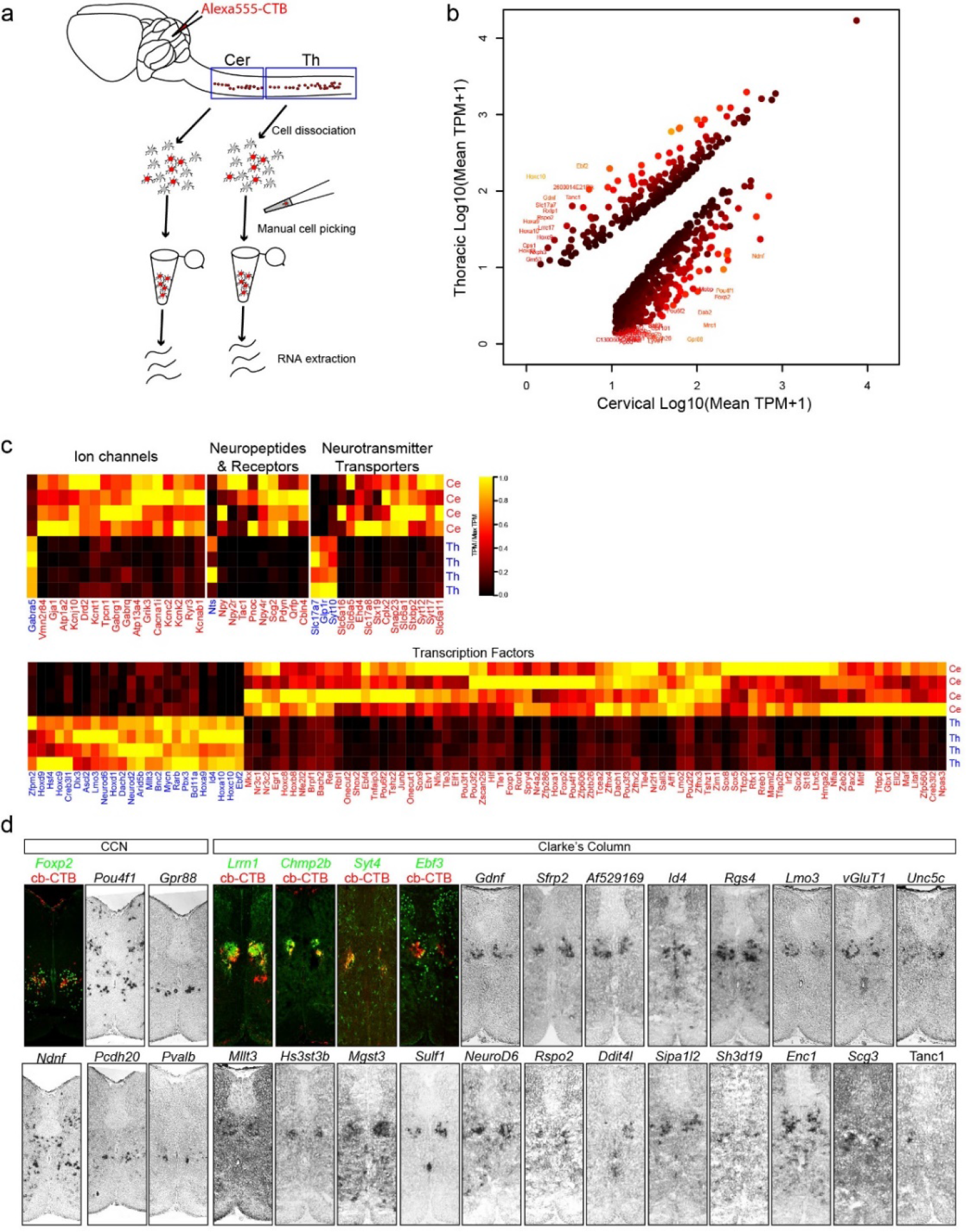
Identification and characterization of CCN and CC molecular markers. (a) Strategy for isolating cervical and thoracic SCTNs for RNAseq. Isolations were performed in quadruplicate at cervical (185, 182, 179, 178 SCTNs between C1-C8) and thoracic (310, 305, 473, 341 SCTNs between T1-T12) levels. (b) Mean expression of differentially expressed genes in cervical (x-axis) and thoracic (y-axis) bulk RNA-seq samples. Genes with differential expression between cervical and thoracic samples with FDR<0.001 (using edgeR), fold-change >2, and mean TPM>10 in either cervical or thoracic samples are shown as dots, colored by FDR value. Genes with fold-change greater than 30 are shown with text labels. (c) Heatmaps showing expression of differentially expressed genes (cervical vs. thoracic, FDR<0.001, fold-change > 2) belonging to major annotated categories. Heatmap colors represent scaled TPM values for each replicate bulk sample. (d) Validation of sequencing data. P6 mouse spinal cord sections were used for validating differential gene expression by in situ hybridization and immunostaining. For identifying SCTNs in immunostaining experiments, Alexa555-CTB labeled spinal cord sections were used.

We further characterized genes differentially expressed between cervical and thoracic SCTNs by performing mRNA in situ hybridization and immunohistochemical analyses (Fig. 2d). We focused on transcription factors, cell adhesion molecules, and genes implicated in neuronal function, as these classes of genes are often selectively and robustly expressed by neuronal subtypes. Most of the cervical enriched genes we identified were expressed in a cluster of neurons located in rostral cervical segments, near the position occupied by CCN neurons. Putative CCN-restricted genes included *Foxp2*, *Pou4f1*, *Gpr88*, *Ndnf*, and *Pcdh20* (Fig. 2d). We confirmed selective expression of *Foxp2* in CCN neurons by performing cerebellar retrograde tracing of SCTNs in conjunction with Foxp2 antibody staining (Fig. 2d, and Supplementary Fig. 2a). This analysis revealed Foxp2 is expressed by labeled SCTNs at rostral cervical levels, but not in caudal cervical or thoracic SCTNs. We also identified a number of genes selective for thoracic CC neurons, including the previously characterized *Gdnf* and *VGlut1* genes^19^. We confirmed SCTN-restricted expression of novel genes, including *Lrrn1*, *Chmp2b*, *Syt4*, and *Ebf3* by performing in situ hybridization or immunohistochemistry in conjunction with cerebellar CTB tracing (Fig. 2d). These genes were expressed selectively by clusters of thoracic neurons, but not in cervical SCTNs, indicating they are selective markers for CC neurons (Supplementary Fig. 2b and data not shown).

### Single cell molecular profiling of SCTNs

To further examine the diversification of SCTNs using genome-wide assays, and to identify smaller subgroups of SCTNs, we performed single cell RNAseq on neurons isolated from rostral cervical, caudal cervical, and rostral thoracic levels. We manually isolated ~100 retrogradely labeled SCTNs from each level and performed scRNAseq. Unsupervised clustering of scRNAseq data identified 8 clusters of neurons (SCT1-8) (Fig. 3a,b, and Supplementary Fig. 3a,b). Two clusters, SCT7 and SCT5, were unique to rostral cervical and rostral thoracic segments and expressed genes indicative of CCN and CC fates, respectively, based on the number and identity of genes that overlapped with our bulk sequencing analyses (Supplementary Fig. 3c). For example, SCT7 expresses *Foxp2* (a CCN marker), while SCT5 expresses *Gdnf* (CC marker). Two clusters SCT2 and SCT3, were present in each of the three segmental levels we analyzed (Fig. 3b), possibly representing Hox-independent populations. Four clusters (SCT1, 4, 6, 8), were present at two levels, with higher representation within a single region. These results potentially identify additional SCTN populations that were likely masked by over-representation of CCN and CC-restricted genes in our bulk sequence analysis.

**Fig. 3.**
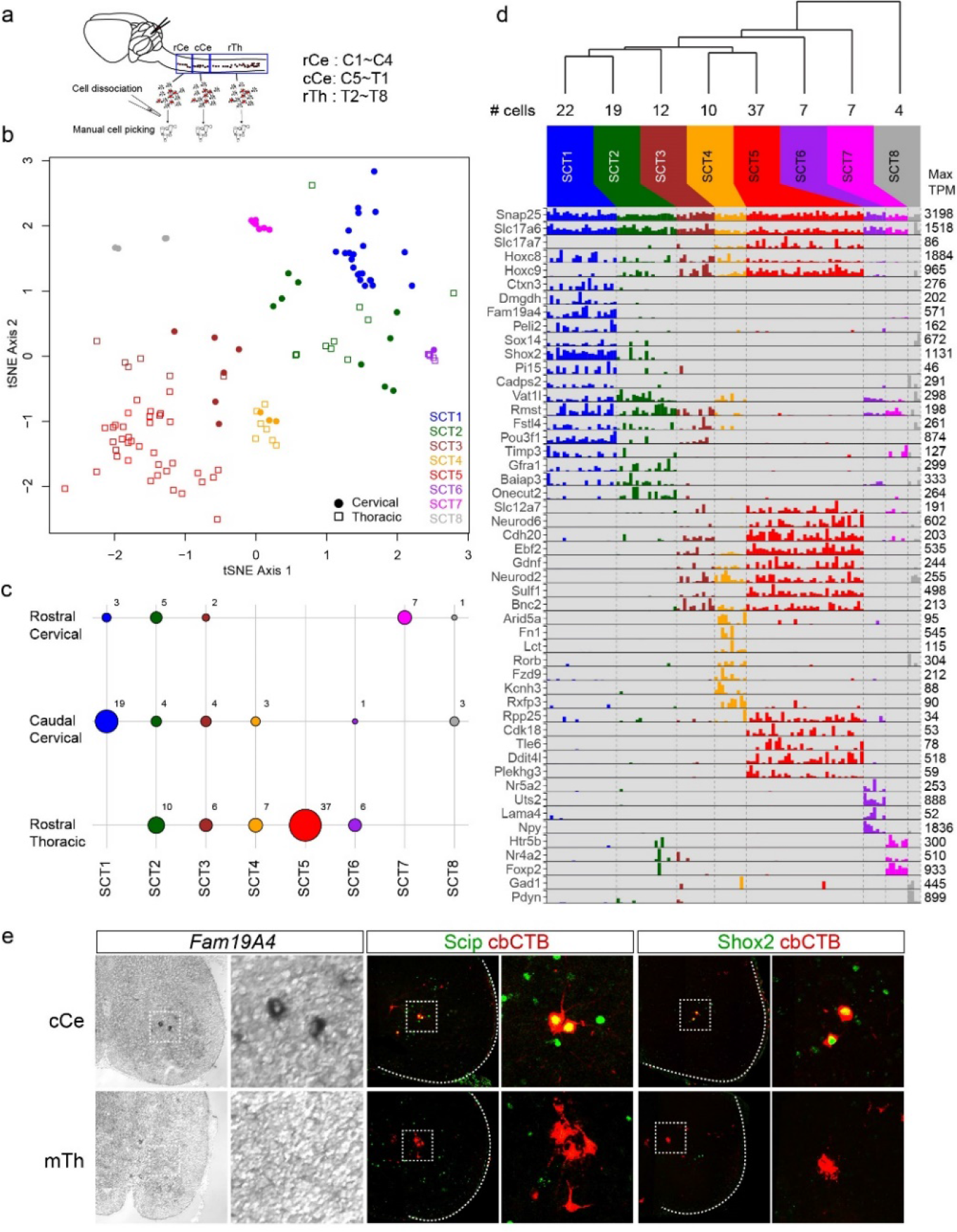
Characterization of SCTN subtypes by scRNAseq. (a) scRNA-seq workflow. Alexa555-CTB was injected into the cerebellum at P4 and cells were harvested from rostral cervical (rCe), caudal cervical (cCe), and rostral thoracic (rTh) spinal cord at P7. SCTNs were manually picked and processed for scRNAseq using a 3’-digital plate-based method. (b) Visualization of putative cell clusters in a t-SNE plot. Cells were clustered as described in the methods (not in t-SNE space), and cluster identities SCT1 through SCT8 are color-coded in the plot. Shapes represent the dissection from which cells were obtained. (c) Dot plot showing the number of cells in each cluster deriving from each segmental dissection. The size of each circle indicated the number of cells in a given cluster from a specific dissection, and the corresponding numbers are indicated to the right of the circles. SCT1 and SCT5 are unique to the cervical and thoracic dissections, respectively. (d) Barplot showing the expression (TPM) values for selected pan-class genes and genes with differential expression across clusters. The hierarchical dendrogram at the top was generated using complete linkage, with the distance metric defined as the Euclidean distance between mean log10(TPM+1) values for each cluster. For each gene, the maximum TPM value is indicated by the number to the right of each row in the bar plot. (e) Expression of *Fam19A4*, Scip, and Shox2 in cCe SCTNs. For Scip and Shox2 analyses, SCTNs were labeled by cerebellar-CTB (cbCTB) retrograde tracing. Labeling of SCTNs with indicated markers were not observed at mid-thoracic (mTh) levels (data not shown).

To determine whether any of our single cell clusters identify additional SCTN types, we chose genes within cluster SCT1 for further analysis. SCT1 neurons derive from caudal cervical segments, possibly representing the LVII SCTN subtype. SCT1 neurons are characterized by elevated expression of *Fam19A4, Shox2*, and *Scip (Pou3f1)* (Fig. 3d). We found that the *Fam19A4* gene was selectively expressed in caudal cervical segments, and marked a small group of spinal neurons (Fig. 3e). We confirmed expression of *Fam19A4* in cervical LVII SCTNs by performing in situ hybridization on spinal cord sections in which SCTNs were labeled through cerebellar retrograde tracing (Supplementary Fig. 3d). Using this approach, we also identified the transcription factors *Shox2* and *Scip* as a selective markers for cervical LVII SCTNs. Although both proteins are expressed throughout the rostrocaudal axis of the spinal cord, we found that Shox2 and Scip were selectively expressed by cerebellar-projecting SCTNs at caudal cervical levels, and labeled the more ventral LVII population (Fig. 3e). Collectively our bulk and single cell RNAseq analyses demonstrate that three SCTN subtypes (CCN, cLVII, and CC) can be molecularly distinguished by differential gene expression.

### Hox protein expression defines SCTN subtypes

What are the mechanisms that determine the diversity and molecular signatures of SCTN subtypes? Because a major difference between SCTNs is their segmental organization, we examined differences in *Hox* gene expression, known determinants of rostrocaudal patterning in the CNS^20^. In vertebrates *Hox* genes are organized in 4 chromosomal clusters, and the position of individual genes within a cluster determines where it is expressed along the rostrocaudal axis. In general, *Hox* genes located at the 3’ end of a cluster are expressed rostrally, while those at the 5’-end are expressed caudally. Analysis of our scRNAseq dataset revealed that cervical and thoracic SCTNs follow this co-linear *Hox* pattern. Rostral cervical SCTNs expressed elevated levels of *Hox4-Hox5* gene paralogs (e.g. *Hoxc4*, *Hoxc5*, and *Hoxa5*), caudal cervical SCTNs expressed *Hox6-Hox8* paralogs (*Hoxc6* and *Hoxc8*), while rostral thoracic SCTNs express *Hox9* genes (*Hoxc9* and *Hoxa9*) (Fig. 4a). In addition, certain *Hox* genes were expressed in multiple segments, suggesting specific combinations of Hox proteins contribute to SCTN specification. For example, *Hoxc8* is detected in both caudal cervical and rostral thoracic SCTNs, while *Hoxc6* is expressed by both rostral and caudal cervical SCTNs (Fig. 4a, Supplementary Fig. 4a).

**Fig. 4.**
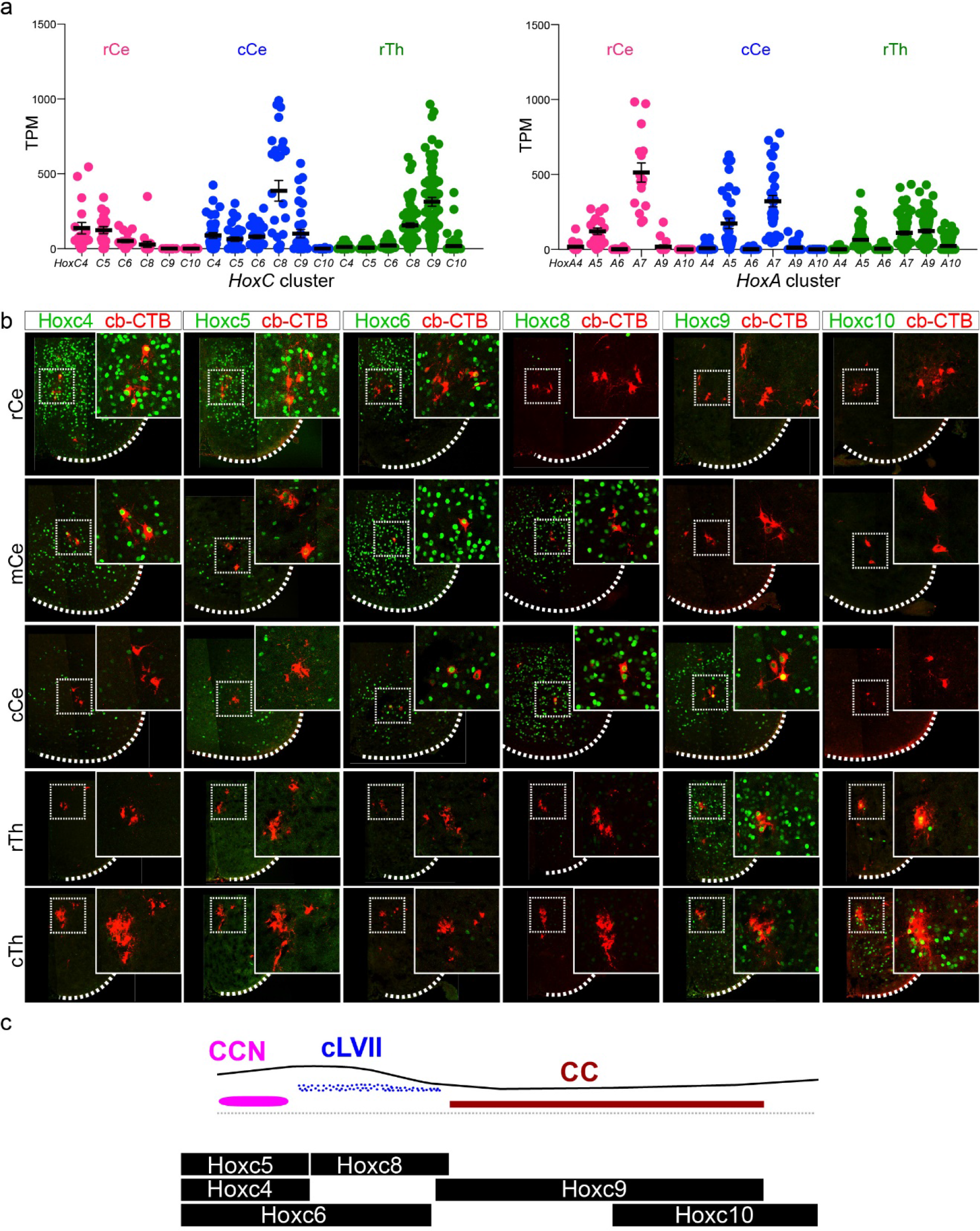
*Hox* expression patterns within SCTN subtypes. (a) Plots of scRNAseq data showing *HoxC* and *HoxA* cluster gene expression levels (TPM) in each segmental region. Only *Hox4-Hox10* paralogs are shown and gene names are abbreviated (e.g. C4 = *Hoxc4*). rCe, 18 cells; cCe, 34 cells; rTh 66 cells. Solid lines indicate mean TPM, error bars +/− S.E.M. (b) Hox protein expression pattern in SCTN subtypes at cervical and thoracic levels. SCTNs were labeled by injection of Alexa555-CTB into the cerebellum at P1 and analyzed using indicated Hox antibodies at P2. (c) Summary of *HoxC* gene expression in cervical and thoracic SCTNs.

We further examined Hox protein expression by performing immunohistochemical analyses in which SCTNs were labeled by cerebellar retrograde tracing at P1. This analysis revealed that cervical CCN neurons express Hoxc4, Hoxc5, low levels of Hoxc6 but lacked Hoxc8 and Hoxc9 expression (Fig. 4b, Supplementary Fig. 4b). Caudal cervical SCTNs express Hoxc6 and Hoxc8, with subsets expressing Hoxc9. Thoracic CC neurons express Hox9 paralogs (Hoxa9, Hoxc9 and Hoxd9) and Hox10 paralogs (Hoxa10 and Hoxc10) (Fig. 4b, Supplementary Fig. 4b,c). Collectively these observations indicate that specific SCTNs populations can be identified by differential expression of Hox proteins, and suggest specific “Hox codes” determine SCTN subtype identity (Fig. 4c).

### *Hox* genes specify SCTN subtype identity

To examine a possible functional role of *Hox* genes in SCTN subtype diversification, we analyzed mice in which specific *Hox* genes are mutated. We first analyzed the effects of mutation of the *Hoxc9* gene, which is normally restricted to thoracic CC neurons. Previous studies have shown that *Hoxc9* is a key determinant of MN subtype identity in thoracic segments, is essential for the generation of preganglionic autonomic MNs, and repression of more anterior *Hox* genes^26,27^. We found that in *Hoxc9* mutants expression of CC-restricted genes was markedly reduced at thoracic levels (Fig. 5a, Supplementary Fig. 5a). Markers normally displaying highly restricted expression in CC neurons, including *Gdnf*, *Syt4*, *Lrrn1*, *Unc5c*, and *Lmo3* were undetectable in thoracic segments of *Hoxc9* mutants (Fig. 5a). Genes which are expressed by CC neurons, but also other spinal populations, such as *Rgs4* and *Id4*, were lost from CC neurons but were preserved in non-SCTN populations (likely representing interneuron populations that do not rely on a specific *Hox* gene or are *Hox*-independent) (Fig. 5a). These observations indicate that *Hoxc9* is necessary for establishing CC-specific gene programs at thoracic levels.

**Fig. 5.**
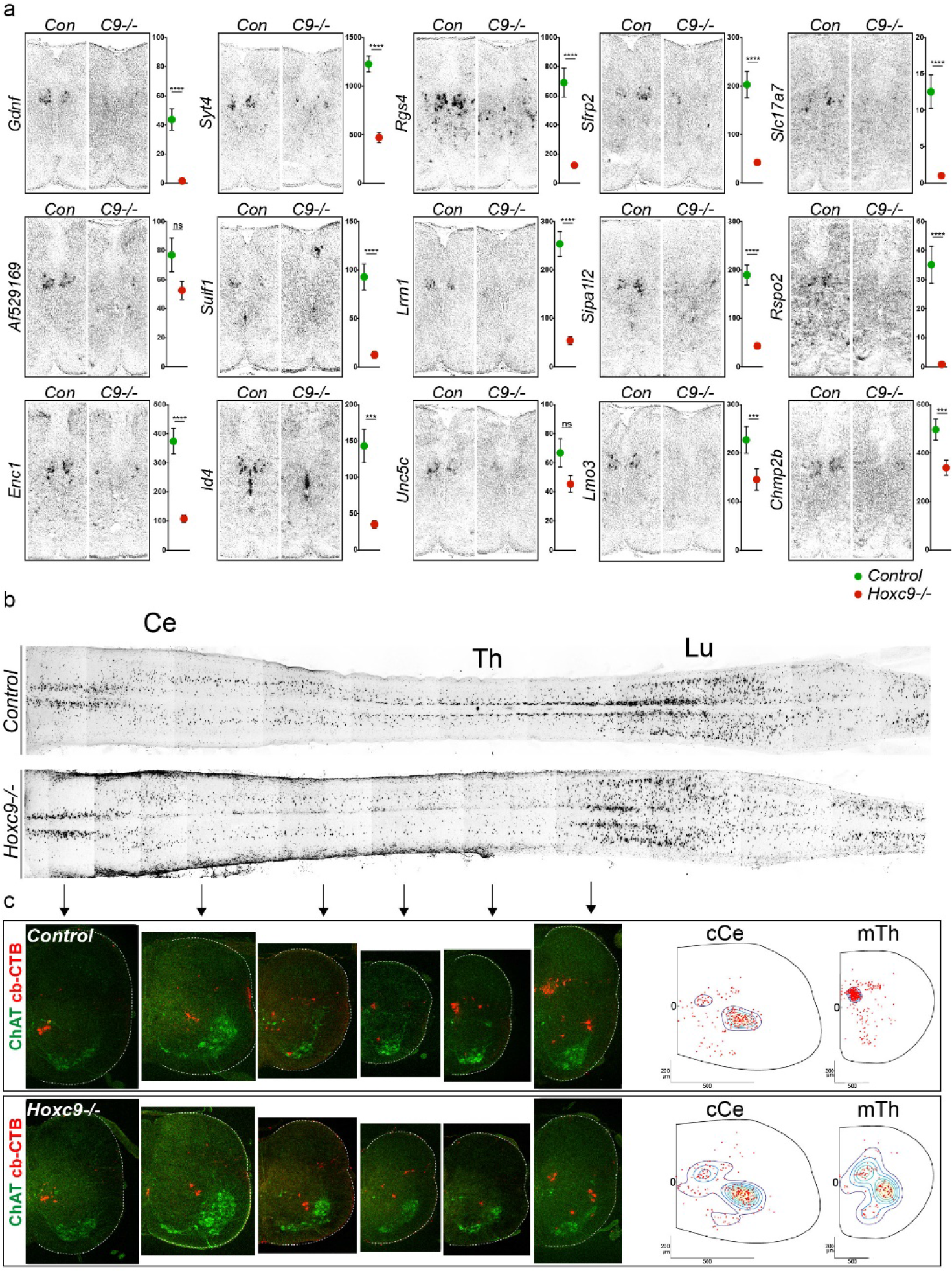
*Hoxc9* is required for Clarke’s column neuron development. (a) In situ hybridization of marker gene expression in control and *Hoxc9* mutants at P6. Graphs on right show scRNAseq data (TPM values) for each gene at thoracic levels in control and *Hoxc9* mutants. P-values of Wilcoxon-Mann-Whitney test: *AF529169*, 0.2153 (ns); *Chmp2b*, 0.0007 (***); *Ddit4l*, <0.0001(****); *Enc1*, <0.0001(****); *Fam19A4*, <0.0001(****); Gdnf, <0.0001(****); Hs3stb1, <0.0001(****); Id4, 0.0006 (***); Lmo3, 0.0001(***); Lrrn1, <0.0001(****); *Pou3f1*, <0.0001(****); *Rgs4*, <0.0001(****); *Rspo2*, <0.0001(****); *Scg3*, <0.0001(****); *Sfrp2*, <0.0001(****); *Sh3d19*, 0.0001(***); *Shox2*, 0.0003(***); *Sipa1l2*, <0.0001(****); *Slc17A7*, <0.0001(****); *Sulf1*, <0.0001(****); *Syt4*, <0.0001(****); *Tanc1*, <0.0001(****); *Unc5c*, 0.1400(ns). P-value of two-tailed Student’s t test: *Mgst3*, <0.0001 (****). (b) Whole mount images of Alexa555-CTB labeled SCTNs in control and *Hoxc9* mutants at P6. (c) Sections of Alexa555-CTB labeled SCTNs at caudal cervical (cCe) and mid-thoracic (mTh) levels. Contour plots are shown on the right. Control cCe, 227cells, n=8 mice; *Hoxc9-/-* cCe, 236 cells, n=11 samples; Control mTh, 340 cells, n=8 mice; *Hoxc9-/-* mTh, 116 cells, n=11mice.

The loss of CC-restricted gene expression in *Hoxc9* mutants suggests *Hox* genes are generally required for deployment of SCTN subtype-specific programs. To further explore this idea, we examined whether additional *Hox* genes are essential during SCTN diversification. We examined the function of *Hoxc8*, which is expressed by caudal cervical LVII SCTNs and characterized by selective expression of *Fam19A4*. We found that in *Hoxc8* mutants expression of *Fam19A4* was lost from the spinal cord (Supplementary Fig. 5b). Interestingly, expression of Scip and Shox2 were retained by some caudal cervical SCTNs (Supplementary Fig. 5c and data not shown), possibly a result of functional compensation by other *Hox* genes. These results indicate that *Hox* genes are essential for the normal specification of SCTN subtypes at cervical and thoracic levels.

The depletion of SCTN markers in *Hox* mutants could be due to the death of these populations at specific segmental levels or a fate switch to an alternate SCTN identity. To assess this at a cellular level, we performed cerebellar retrograde tracing to determine whether any SCTNs are generated in thoracic segments of *Hoxc9* mutants. We injected CTB into the cerebellum of *Hoxc9* mutants and mapped the position of labeled SCTNs. We found that in *Hoxc9* mutants the dorsomedial population of CC neurons is no longer labeled in thoracic segments, with only a small population present at rostral lumbar levels (Fig. 5b,c). SCTNs were labeled in thoracic segments but were scattered and resided in a position similar to those of caudal cervical LVII types (Fig. 5b,c). In contrast, labeling of SCTNs at caudal cervical levels was similar between control and *Hoxc9* mutants, indicating a selective function of *Hoxc9* in thoracic SCTNs. These results indicate that in the absence of *Hoxc9* thoracic SCTNs acquire the settling characteristics of cervical LVII SCTNs.

### Clarke’s column is transformed to a cervical SCTN identity in *Hoxc9* mutants

The acquisition of LVII neuron characteristics by thoracic SCTNs suggests a possible identity transformation in *Hoxc9* mutants. To examine a potential fate conversion at a molecular level, we assessed global changes in the transcriptomes of SCTNs in absence of *Hoxc9* function. We compared scRNAseq profiles from rostral thoracic SCTNs isolated from control and *Hoxc9* mutants, and compared these with control rostral and caudal cervical SCTN populations (Fig. 6a). We found that rostral thoracic SCTNs lacking *Hoxc9* failed to form the CC cluster (SCT5), and the transcript levels of CC-restricted genes were markedly reduced (Fig. 5a). The molecular profile of many thoracic SCTNs in *Hoxc9* mutants matched those of caudal LVII SCTNs (SCT1) (Fig. 6b,c). Upregulated genes in *Hoxc9* mutants included those we identified in our scRNAseq of control caudal cervical SCTNs, including *Hoxc8, Fam19A4*, *Scip (Pou3f1)*, and *Shox2* (Fig. 6d). SCT3, which is normally found at all segmental levels, was still present in thoracic SCTNs of *Hoxc9* mutants (Fig. 6b), consistent with a specification program that is independent of a specific *Hox* gene. These results indicate that in absence of *Hoxc9* thoracic CC neurons acquire the molecular profile of cervical SCTNs.

**Fig. 6.**
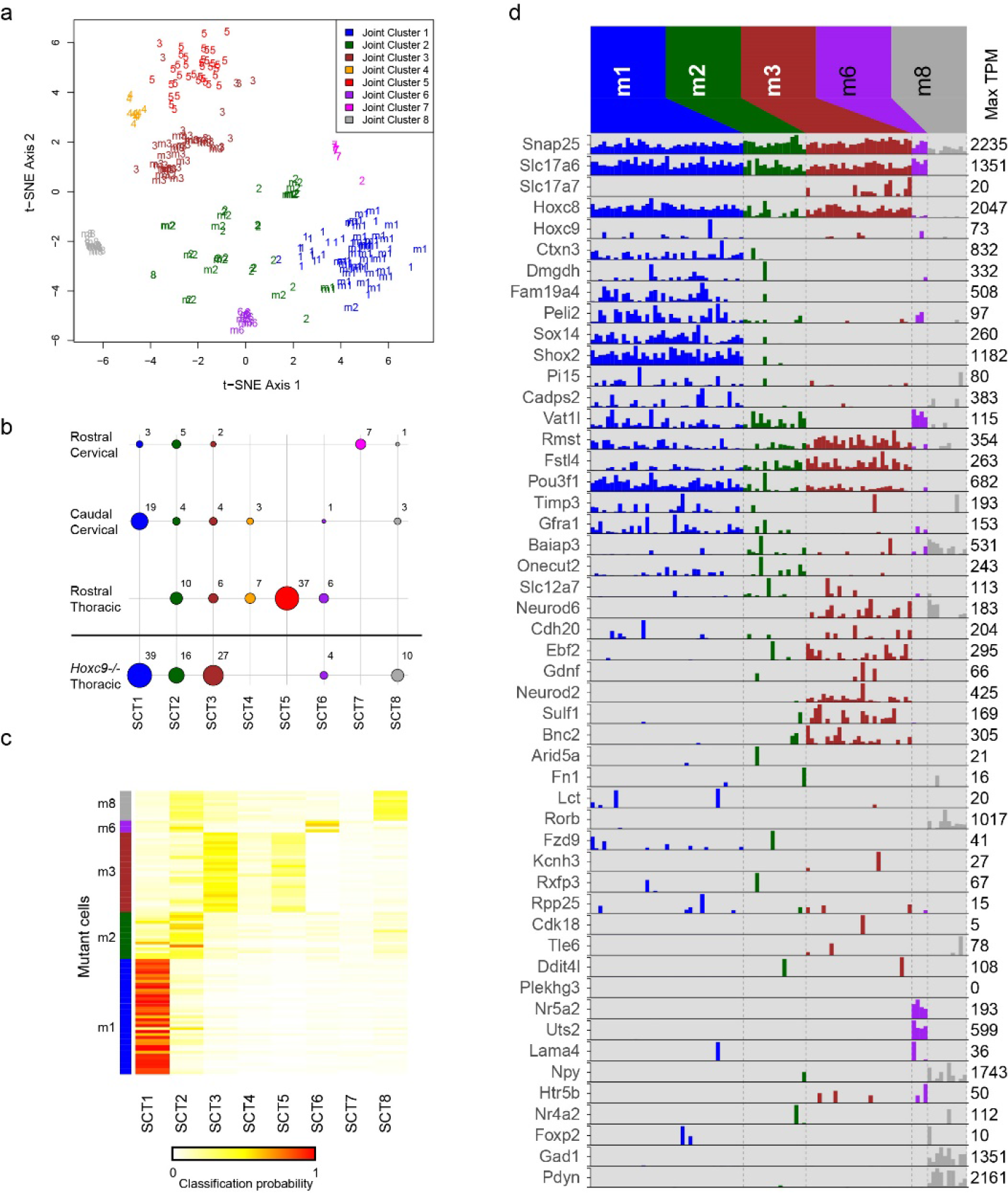
Single cell RNA sequencing of thoracic SCTNs in *Hoxc9* mutants. (a) t-SNE visualization of putative joint and separate cell clusters using all cells from both control and *Hoxc9* mutants. Cells were clustered in three sets: 1) control only (as represented in Fig. 3), which are labeled as 1-8 on the plot, corresponding to SCT1 through SCT8, 2) mutant-only, which are labeled as m1, m2, m3, m6, m8 on the plot, and 3) both control and mutant cells; these joint clusters are color-coded on the plot. In general, the joint clusters agree with the independent clustering of control-only and mutant-only cells, and suggest the correspondence across the two sets of cells. For example, joint cluster 1 (blue) contains cells mostly from control SCT1 (1) and mutant cluster 1 (m1), while joint cluster 3 (brown) contains cells mostly from control SCT3 (3) and mutant cluster 3 (m3). (b) As in Fig. 3C, dot plot representing the number of cells in each cluster originating from each control and mutant dissection. For the mutant, each cluster was assigned to its corresponding control SCT cluster, based on the joint clustering shown in panel A. The numbers for the control dissections are the same as in Fig. 3C. None of the *Hoxc9* mutant cells corresponded to SCT4, SCT5, and SCT7 clusters from the control. (c) Alternative approach to assign *Hoxc9* mutant cells to control clusters. The heatmap shows the classification probabilities for each mutant cell (row) using a random forest classifier trained on the 8 control cluster identities (columns). The colorbar on the left indicates the mutant cell cluster identity (m1, m2, m3, m6, m8). The overall classification closely resembles the result from the joint clustering shown in panel A; for example, cells from m1 have high classification scores for control SCT1, whereas cells from m3 tend to be most strongly assigned to SCT3. (d) As in Fig. 3D, barplot showing expression (TPM) values for selected genes in the cell clusters derived from *Hoxc9* mutant SCTNs.

To further characterize the transformation of CC neurons in *Hoxc9* mutants, we examined whether genes normally enriched in caudal cervical SCTNs are derepressed at thoracic levels. Consistent with our scRNAseq data, as well as previous studies on *Hoxc9* function in spinal MNs, Hoxc8 protein was derepressed in thoracic SCTNs of *Hoxc9* mutants (Fig. 7a,b). Retrograde tracing of SCTNs in *Hoxc9* mutants confirmed that labeled thoracic SCTNs ectopically express Hoxc8 (Fig. 7a). In addition, expression of Hoxc10 was lost from SCTNs at thoracic levels (Supplementary Fig. 7a). We also analyzed expression of Scip and Shox2 proteins, two markers enriched in caudal cervical SCTNs. The number of thoracic SCTNs expressing Scip and Shox2 were markedly increased in *Hoxc9* mutants (Fig. 7c-f). In addition, *Fam19A4*, a selective marker for caudal cervical SCTNs, was ectopically expressed in *Hoxc9* mutants (Fig. 7g,h). The transformation of CC neurons to a cervical LVII fate was also observed in rostral thoracic segments of *Nestin::Cre;Hoxc9 flox/flox* mice, indicating this identity switch is due to a neural-specific function of *Hoxc9*, and not general defects in early rostrocaudal patterning (Supplementary Fig. 7b-d). These results indicate that in the absence of *Hoxc9*, thoracic SCTNs acquire both the anatomical settling position and molecular identity of caudal cervical LVII neurons.

**Fig. 7.**
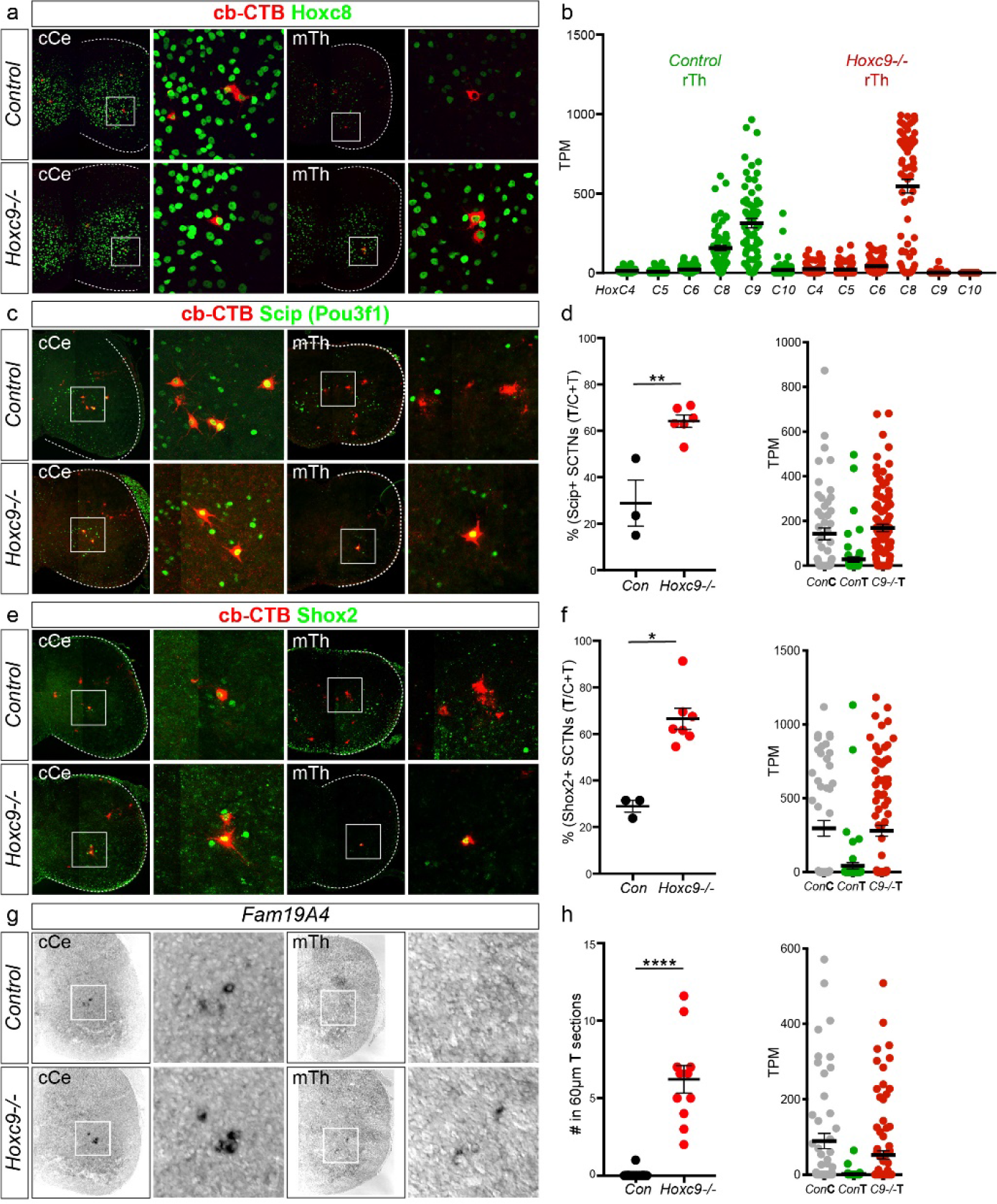
Thoracic SCTNs are transformed to a cLVII identity in *Hoxc9* mutants. (a) Immunostaining of Hoxc8 in retrogradely labeled SCTNs in control and *Hoxc9* mutants in cCe and mTh segments. Thoracic SCTNs express Hoxc8 in *Hoxc9* mutants. (b) Quantification of *Hox* gene expression in single cells from control and *Hoxc9* mutant thoracic regions. Solid lines indicate means, error bars S.E.M. (c) Ectopic expression of Scip (Pou3f1) in thoracic SCTNs of *Hoxc9* mutants. (d) Quantification of Scip^+^ SCTN number and TPM values in single cells from control cervical (ConC), control thoracic (ConT) and *Hoxc9* mutant thoracic (C9−/−T) regions. For the Scip^+^ cell quantification cells were counted in regions belong to LVII group according to contour plot in Fig. 1. Two tailed Student’s t test, p=0.0023 (**); Con, n=3, 81cells (Ce, 57; Th, 24); *Hoxc9−/−*, n=6, 156cells (Ce, 54; Th,102). (e) Ectopic expression of Shox2 in thoracic SCTNs of *Hoxc9* mutants. (f) Quantification of Shox2^+^ SCTN number and TPM scRNAseq values of ConC, ConT, and C9−/−T regions. Wilcoxon-Mann-Whitney test, p=0.0167 (*); Con, n=3, 91cells (Ce, 64; Th, 27)*; Hoxc9−/−*, n=7, 179 cells (Ce, 59; Th, 120). For the Shox2^+^ cell quantification cells were counted in regions belong to LVII group according to contour plot in Fig. 1. (g) Ectopic expression of *Fam19A4* in thoracic sections of *Hoxc9* mutants. (h) Quantification of *Fam19A4*+ cells and TPM single cell values of ConC, ConT, and C9−/−T regions. Wilcoxon-Mann-Whitney test, p<0.0001 (****); Con, n=4 (P6, 2; E15.5, 2; 21sections); *Hoxc9−/−*, n=2 (P6, 1; E15.5, 1; 11sections).

We also asked whether loss of *Hoxc8*, which is required for acquisition of cervical LVII SCTN molecular features, leads to a similar transformation in identity. In *Hoxc8* mutants, Hoxc4 and Hoxc5 were derepressed in caudal cervical segments (Supplementary Fig. 5d). In addition retrograde tracing from the cerebellum indicated that labeled caudal cervical SCTNs ectopically express *Hoxc4*, suggesting a fate switch to a more rostral identity (Supplementary Fig. 5e). However, analysis of CCN marker expression, including *Foxp2* and *Gpr88*, failed to reveal a transformation in SCTN identity (data not shown). The absence of a complete fate transformation in *Hoxc8* mutants is likely due to presence of additional *Hox* genes in caudal cervical segments, which could lead to an ambiguous Hox code.

### Transformation of SCTN identity disrupts spinocerebellar circuitry

Our results indicate that in the absence of *Hoxc9* thoracic SCTNs are converted to a cervical LVII SCTN molecular identity. We examined whether this switch in transcriptional profile is accompanied by changes in the connectivity between SCTNs, pSNs, and the cerebellum. We first assessed whether the loss of CC identity in *Hoxc9* mutants affects innervation of the cerebellum by SCTN axons. Because the number of thoracic SCTNs is markedly reduced in *Hoxc9* mutants (Fig. 5b), we tested whether there is an overall loss of innervation. To label precerebellar SCTN axons, we injected an AAV virus expressing GFP under the *synaptophysin* promoter into rostral cervical and thoracic segments, and examined axonal termination patterns (Supplementary Fig. 8a). In control animals, injections into rostral cervical segments (containing CCN neurons) labeled axons that terminate in lobules 2, 3, 4/5, and 9. Injection of viral tracer into thoracic segments exhibited denser cerebellar innervation that terminated in lobules 2, 3, 4/5, 8 and 9. In *Hoxc9* mutants the overall density of projections from thoracic segments to the cerebellum was markedly reduced, despite levels of GFP expression in the spinal cord that were similar to controls (Supplementary Fig. 8a,b). These observations indicate that loss of *Hoxc9* erodes the normal profile of connectivity between thoracic SCTNs and the cerebellum.

Caudal cervical LVII SCTNs receive input from pSNs that target forelimb muscle. If the transformation of CC neurons to a caudal cervical LVII identity switches their connectivity, they might now receive ectopic inputs from the central afferents of forelimb pSNs. We therefore examined whether ectopic thoracic LVII SCTNs receive forelimb muscle input. We injected CTB into forelimb muscles of control and *Hoxc9* mutant animals, while in parallel tracing SCTNs through injection of HRP into the cerebellum. Synapses between CTB-traced proprioceptors onto HRP+ SCTNs was determined by costaining with VGlut1. The number of ectopic synapses from limb proprioceptors to thoracic SCTNs was markedly increased in *Hoxc9* mutants (Fig. 8a-c). These results indicate that the transformed SCTNs in *Hoxc9* mutants receive presynaptic inputs appropriate for their switch in identity. Collectively these results show that *Hox* genes are essential for the subtype diversification and connectivity of spinocerebellar circuits.

**Fig. 8.**
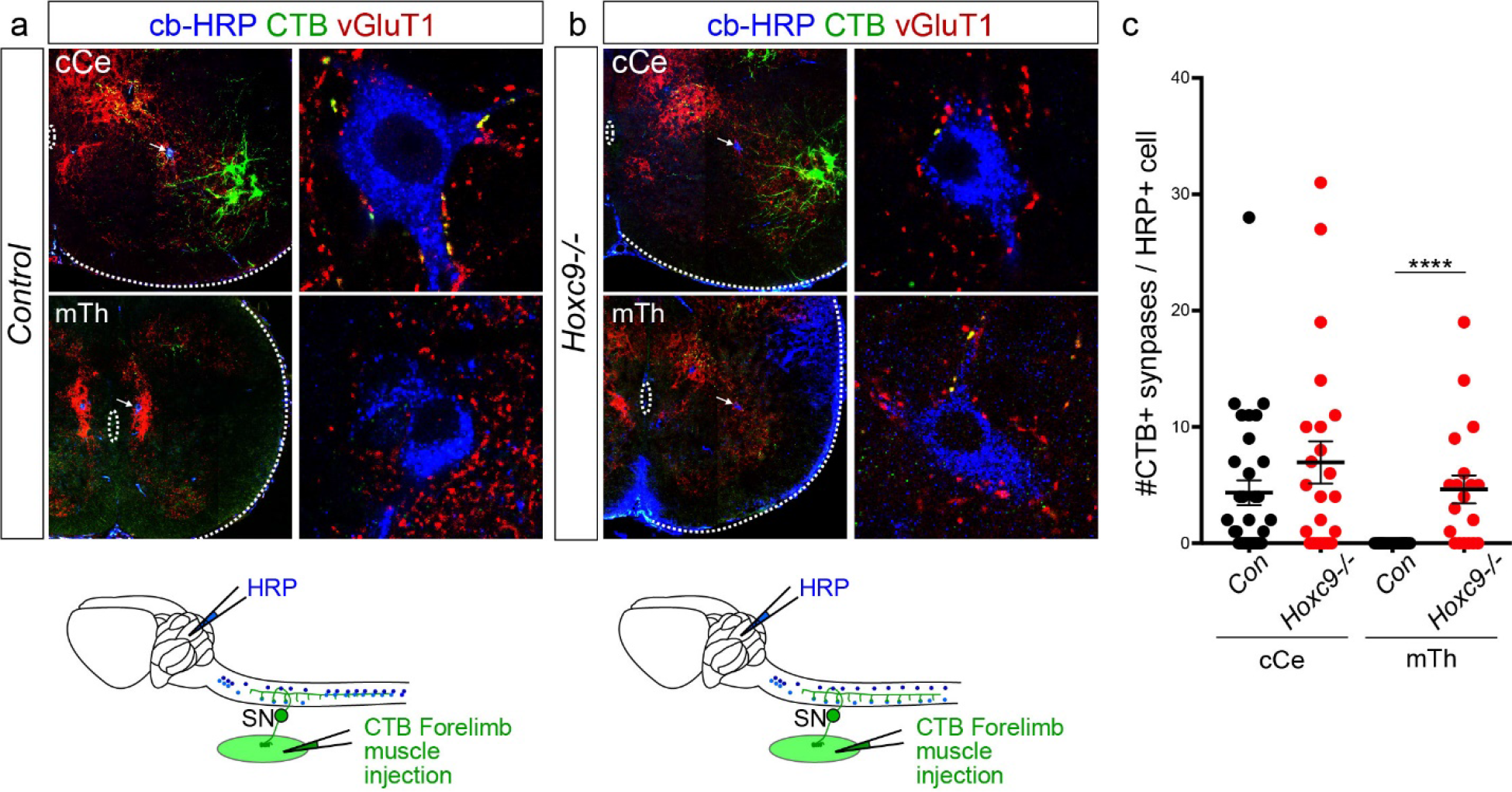
Forelimb pSNs synapse with thoracic SCTNs in *Hoxc9* mutants. (a-b) Immunostaining of cCe and mTh spinal sections of control and *Hoxc9* mutants. SCTNs were labeled via injection of HRP into the cerebellum, and forelimb pSN central afferents were traced by CTB intramuscular injection. (c) Quantification of synapses between forelimb pSNs and SCTNs. For control cCe, mutant cCe, control mTh, and mutant mTh quantification synapses were counted in cells belonging to LVII and CC group according to contour plot in Fig. 1. Controls: cCe, 32 cells; mTh, 24 cells; n=4 mice (3 triceps, 1 forelimb distal flexor muscles). *Hoxc9* mutant: cCe, 23 cells; mTh, 19 cells; n=4 mice (3 triceps, 1 forelimb distal flexor muscles). Wilcoxon-Mann-Whitney test, p<0.0001 (****).

## Discussion

Control of movement depends on accurate reporting of muscle and joint contractile status from proprioceptive sensory neurons to the CNS. Proprioceptive information is relayed to the cerebellum through diverse SCTN subtypes, but the molecular logic by which SCTN identity and connectivity is achieved is largely unknown. By combining single-cell molecular profiling and genetic analyses, we have identified a Hox-dependent genetic program essential for the diversification and synaptic specificity of SCTNs that relay proprioceptive sensory information from limb and axial muscle to the cerebellum. Our findings indicate that the same developmental mechanisms used to generate the diversity of spinal MNs are essential for specifying subtypes of sensory-relay interneurons. These results suggest a general mechanism through which a single large family of transcription factors establishes the diversity of multiple neuronal classes.

### Molecular and Anatomical Diversity of SCTNs

Using genome-wide interrogation of SCTN subtypes generated at cervical and thoracic levels, we identified molecular signatures which distinguish CCN, cLVII, and CC neurons, three major SCTN subtypes that relay proprioceptive information from neck, forelimb, and hindlimb muscles, respectively. Our scRNAseq analysis identified 8 clusters of neurons, each likely representing a specific SCTN subtype. We found that three of these clusters, SCT1, SCT5, and SCT7 represent cLVII, CC, and CCN subtypes and constitute the majority of SCTN populations generated at cervical and thoracic levels. The additional 5 clusters we identified could represent smaller subtypes of SCTNs, such as the more scattered populations normally observed at multiple segmental levels. These populations likely encompass SCTN lineages derived from spinal progenitors expressing the transcription factor *Atoh1*^28,29^, which includes a population recently shown to define a distinct group of non-Clarke’s column SCTNs^30^.

### Role of *Hox* Genes in Determining SCTN Organization and Subtype-Specific Features

Our studies indicate that Hox transcription factors play critical roles in specifying SCTN subtype identity at cervical and thoracic levels. We found that SCTN subtypes can be defined by expression of specific Hox transcription factors. CCN neurons express *Hox5* paralogs, cLVII neurons express *Hoxc8*, while CC neurons express *Hox9* and *Hox10* genes. Mutation in the thoracic *Hoxc9* gene leads to a loss of CC-specific molecular programs, while mutation in *Hoxc8* erodes the molecular specification of cLVII neurons. In the absence of *Hoxc9*, all molecular features of thoracic CC neuron are depleted, with only lumbar-level expression of these genes being maintained. The preservation of CC identity at lumbar levels suggests multiple *Hox* genes are involved in specifying CC features, which likely include additional genes in the *Hox9* and *Hox10* paralog groups. Similarly, the regulation of rostral cervical CCN-restricted determinants likely requires the activities of multiple *Hox5* paralogs.

Recent studies suggest that molecular programs acting along the rostrocaudal axis play key roles in establishing subtype-specific features of spinal interneuron classes. Both V1 and V2a interneuron classes are generated from a single progenitor domain, but give rise to dozens of molecularly distinct subtypes, which can be defined through differences in settling position, connectivity, and transcription factor gene expression^21,22,31,32^. While studies of V1 interneurons have demonstrated an important role of *Hox* genes in patterning transcription factor expression^21^, the identities of their subtype-specific targets are unclear. We found that in the absence of *Hoxc9*, expression of dozens of CC-restricted markers are markedly reduced. In both *Hoxc8* and *Hoxc9* mutants, the more rostrally expressed *Hox* genes are derepressed, similar to the boundary-maintenance function of Hox proteins observed in MNs^20^. This leads to either a transformation in SCTN fate as in *Hoxc9* mutants, or a disruption in normal specification programs, as seen in *Hoxc8* mutants. These findings suggest that similar to MNs, the diversification of spinal interneuron classes relies on Hox-dependent transcriptional networks to both activate and repress repertoires of subtype specific genes.

### Establishing synaptic specificity in proprioceptive sensory circuits

Our studies provide insights into developmental mechanisms through which proprioceptive circuits are assembled. After entering the spinal cord, proprioceptive sensory neurons establish highly specific connections to diverse classes of post-synaptic targets. The best studied pSN connections are those established with MNs^9,33^. Each pSN forms a specific connection to the MN pool that targets the same or functionally related muscle, while avoiding MNs targeting antagonistic muscles. These connections are highly selective, such that a single pSN targets each of the ~50-100 MNs within the entire pool that supplies the same peripheral muscle^34^.

How the striking synaptic specificity between pSNs and their central synaptic targets is achieved is poorly understood, but appears to involve both genetic and activity-dependent processes^35,36^. Mutations in genes involved in pSN fate determination, such as the transcription factors *Er81* or *Runx3*, leads to widespread defects in the connectivity and survival of pSNs^37–39^. Recent studies indicate that postsynaptic, target-derived features shape the specificity between pSN and MNs^40,41^. For example, transforming the identity of thoracic MNs to a limb-level fate, through deletion of the *Hoxc9*, causes limb-derived pSNs to target MNs present at thoracic levels^42^. These observations indicate that subtype identity of postsynaptic targets plays an instructive role in determining connectivity with pSNs.

In contrast to the specific point-to-point central connectivity between pSNs and MNs innervating the same muscle, connections between pSNs and SCTNs appear to be less restricted. Individual neurons within Clarke’s column receive direct and indirect proprioceptive inputs from multiple, often functionally antagonistic muscle groups^17,18^. Nevertheless, the specificity of inputs from pSNs to SCTNs could be restricted by the identity of the muscle source (e.g. forelimb versus hindlimb). We found that transformation of SCTNs identities leads to changes in their pre- and post-synaptic target specificity. In *Hoxc9* mutants, forelimb pSNs establish connections to the cLVII neurons ectopically generated in thoracic segments. These results parallel the circuit alterations between pSNs and MNs observed in *Hoxc9* mutants, where forelimb pSNs synapse with the ectopically generated thoracic lateral motor column MNs^42^. It appears therefore that as pSN central afferents enter the spinal cord, target specificity is shaped by recognition of molecular differences in the subtypes of neurons they encounter.

A notable feature of Clarke’s column is an absence of registry between its segmental position and the location of the pSNs from it receives direct input. Most CC neurons are located at thoracic levels, while hindlimb pSNs reside in lumbar segments. This positional mismatch could be attributed to a change in CC function during vertebrate evolution. One possibility is that SCTNs with CC-like molecular features were initially used for relaying proprioceptive information from non-limb axial muscle. In fish, reptiles, and amphibians axial muscles play prominent roles in coordinating locomotor behaviors and likely required spinocerebellar pathways during motor control. The appearance of paired appendages might have attenuated the importance of axial proprioception, while hindlimb pSNs co-opted the existing thoracic system for limb-based locomotion. The *Hoxc9* gene appears to exert an important role in maintaining this ancestral SCTN genetic program, in part, by suppressing expression of *Hox* genes associated with forelimb-level spinal neurons, and enabling thoracic expression of hindlimb-associated *Hox10* genes.

The organization of SCTNs into columnar groups was likely a later mammalian innovation, as cervical and thoracic SCTNs of amphibians and reptiles do not appear to form longitudinal clusters^43,44^. SCTN columnar organization may have evolved in mammals to facilitate additional layers of interconnectivity, such as those with descending motor pathways or between different types of sensory afferents. An exciting future direction will be to identify the class-specific factors that cooperate with Hox proteins, and their downstream effectors which construct neuronal columns in mammalian spinal cord.

Studies in humans and animal models indicate that loss of muscle-derived sensory information does not prohibit the ability of spinal circuits to generate basic motor output, but is essential for adaptive behaviors and motor learning. The relative contributions of proprioceptive input to local spinal networks versus ascending pathways in motor control are unclear. Mice that lack muscle spindles or pSNs display defects in locomotor coordination^7,37,45^, but whether this is due to alteration in pSN connections to spinal neurons, spinocerebellar circuits, or both is unknown. The identification of selective molecular features of SCTNs should provide means to ascertain the relative contributions of spinal and supraspinal proprioceptive pathways to motor control. These studies may provide insights into how sensory-motor information is integrated at the level of the spinal cord, as well basic insights relevant to the study of spinocerebellar ataxias.

## Online Methods

### Mouse genetics

Animal work was approved by the Institutional Animal Care and Use Committee of the NYU School of Medicine in accordance with NIH guidelines. Mouse lines used were: *Hoxc9 flox*^42^, *Hoxc9-/-* ^26^, *Hoxc8-/-*^46^, *Nestin::Cre* (The Jackson Laboratory #003771), FVB (#207, Charles River Lab).

### Immunohistochemistry

Antibodies against Hox proteins have been previously described^27,47^. Additional antibodies used were the following.

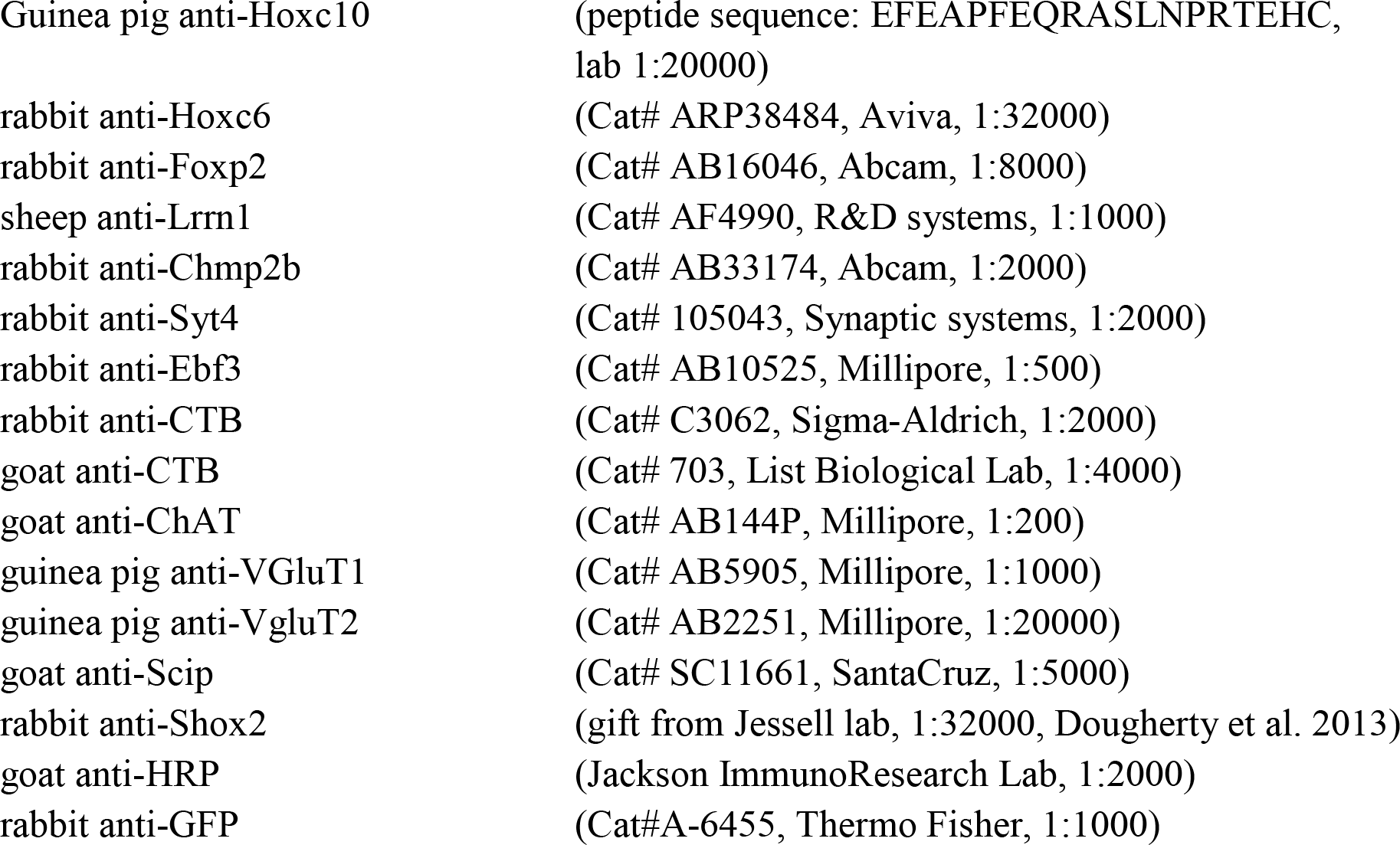

### Secondary antibodies

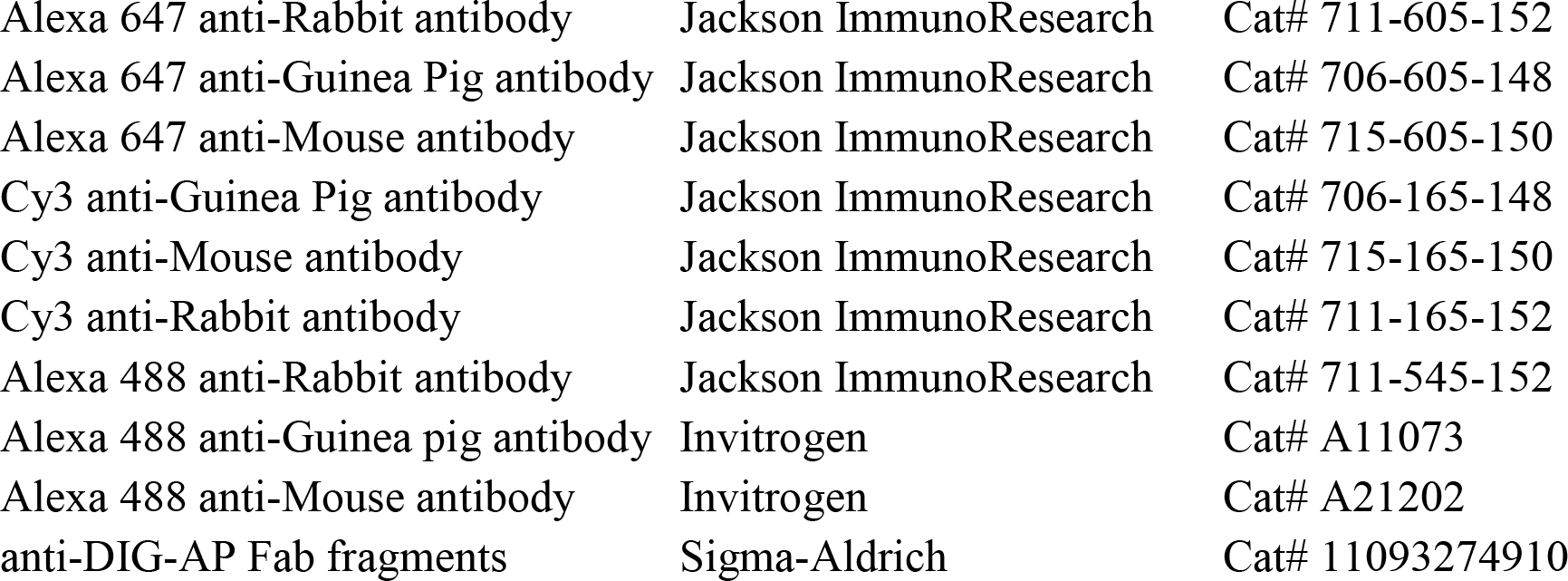

*Detailed protocols for histology are available on the Dasen lab website (http://www.med.nyu.edu/dasenlab/:).*

### In situ hybridization

In situ hybridization of tissue sections was performed as previously described using DIG labeled probes^48^.

### SCTNs labeling, isolation, and RNA sequencing

SCTNs were labeled by injecting CTB (Alexa555 conjugated form, 1µg/µl in PBS, Cat# C34775, Invitrogen) into the cerebellum using NanojetII(Cat# 3-000-204, Drummond Scientific Company) at P4 and examined at P6-P7. Labeled SCTNs were collected manually as described^49^with some modifications: before pronase incubation meninges were removed as much as possible and 150-300μm transverse spinal cords slices were generated using a razor blade.

#### Bulk RNA sequencing

Slices were incubated in ACSF (126mM NaCl, 3mM KCl, 1.25mM NaH2PO4, 20mM NaHCO3, 20mM D-Glucose, 2mM CaCl2, 2mM MgCl2 w/ pronase) for 50min. During cell collection for bulk sequencing, neuronal activity blockers were not included in ACSF. Sorted cells were transferred to tubes containing 50μl Picopure RNA extraction buffer. RNAs were extracted and spiked in ERCCs. Sequencing libraries were prepared using NuGEN SPIA library prep kit. Quadruplicates of pooled samples were used for bulk sequencing: Cervical (C1-C8;185, 182, 179, 178cells)/Thoracic (T1-T12, 310, 305, 473, 341 cells)

#### Single cell RNA sequencing (protocol designed by Janelia Research Campus)

During cell collection for single cell RNA sequencing, neuronal activity blockers (TTX, APV, and DNQX) were included in ACSF as described ^49^. Slices were incubated in ACSF (w/ pronase) for 50min. After dissociation of labeled cells, each cell was transferred to 0.2 ml PCR 8-tube strip (1402-4700, USA Scientific) containing 3 ul lysis buffer (0.2% Triton X-100 (Cat#T8787-100ML, Sigma Aldrich) in Nuclease-free water (Cat#AM9937, Ambion) with 0.1 U/ul RNase inhibitor (Cat#30281-1, Lucigen). During cell transfer, 0.1-0.2 ul ACSF cocktail was transferred to the collection tube. Each 8-tube strip of cells was flash frozen on dry ice and kept at −80ºC until sequencing experiment was performed. Number of cells used in single cell sequencing: MRT(T2-T8), 125cells; CRT(T2-T8), 78cells; CCC(C5-T1), 53cells; CRC(C1-C4), 23cells. RNAseq data is available through GEO (accession in progress)

### SCTNs and sensory terminal labeling

SCTNs were labeled by injecting HRP (20%, 100mg HRP (Cat# 814 407, Roche) dissolved in 1% Lysophosphatidyl choline (Cat# L4129, Sigma Aldrich) PBS) into the cerebellum and muscle sensory terminals were labeled by CTB ((2% CTB; Sigma-Aldrich, Cat# C9903; buffer was changed to PBS using Amicon Ultra-0.5 Centrifual filter units, Cat# UFC503008, Millipore) injection into the muscle at P4. Samples were perfused (4% PFA), saturated with sucrose (30%), and cryosectioned at 30um. Signals were examined at P6 using immunohistochemistry.

### AAV injection into the spinal cord

Retrograde AAV variant (0.5μl, AAV-SL1-synGFP, gift from Janelia Research Campus) was injected into the spinal cord at P1 using NanojetII and examined at P6. Injected samples were perfused (4%PFA), saturated with sucrose (30%), and cryosectioned at 40um.

### Image acquisition

Zeiss confocal microscope (LSM700, 20X dry or 63X oil objective lenses) was used for acquiring images. Images were processed in Fiji and Photoshop.

### Contour plots

Images were fit to the representative spinal cord sections using the landmark correspondence plugin in ImageJ. X-Y coordinates were acquired in ImageJ. Isoline plots were generated from X-Y scatter plots using Bivariant Kernel Density Estimation function (gkde2) with default setups in MATLAB. Nine isolines (from yellow to blue) were generated by default: yellow line, most dense region; blue line, least dense region.

### Quantification and statistical analysis

Statistical analysis was performed using Prism 7 software. Normality test was performed before sample comparison (Shapiro-Wilk normality test or D’Agostino & Pearson normality test). If samples were met normality criteria, samples were compared using two-tailed Student’s t test; if not, non-parametric (Wilcoxon–Mann–Whitney) tests were used.

## Acknowledgements

We thank Karel Liem for providing *Hoxc9* mutants, Helen Lai for sharing unpublished results, and Thomas Jessell for advice and guidance. We thank Kristen D’Elia, Sara Fenstermacher, Britton Sauerbrei, and David Schoppik for discussion and feedback on the paper. This work was supported by NIH R01 NS097550 and NS062822 from NINDS to J.S.D. and funding from HHMI to A.H.

## Author Contributions

M.B. performed all of the mouse genetic experiments, histological analyses, and isolated SCTNs for RNAseq. V.M. analyzed the bulk and scRNAseq data. All authors read, edited and approved the final manuscript.

## Competing Interests

The authors declare no competing financial interests.

---

**Supplementary Fig. 1.**
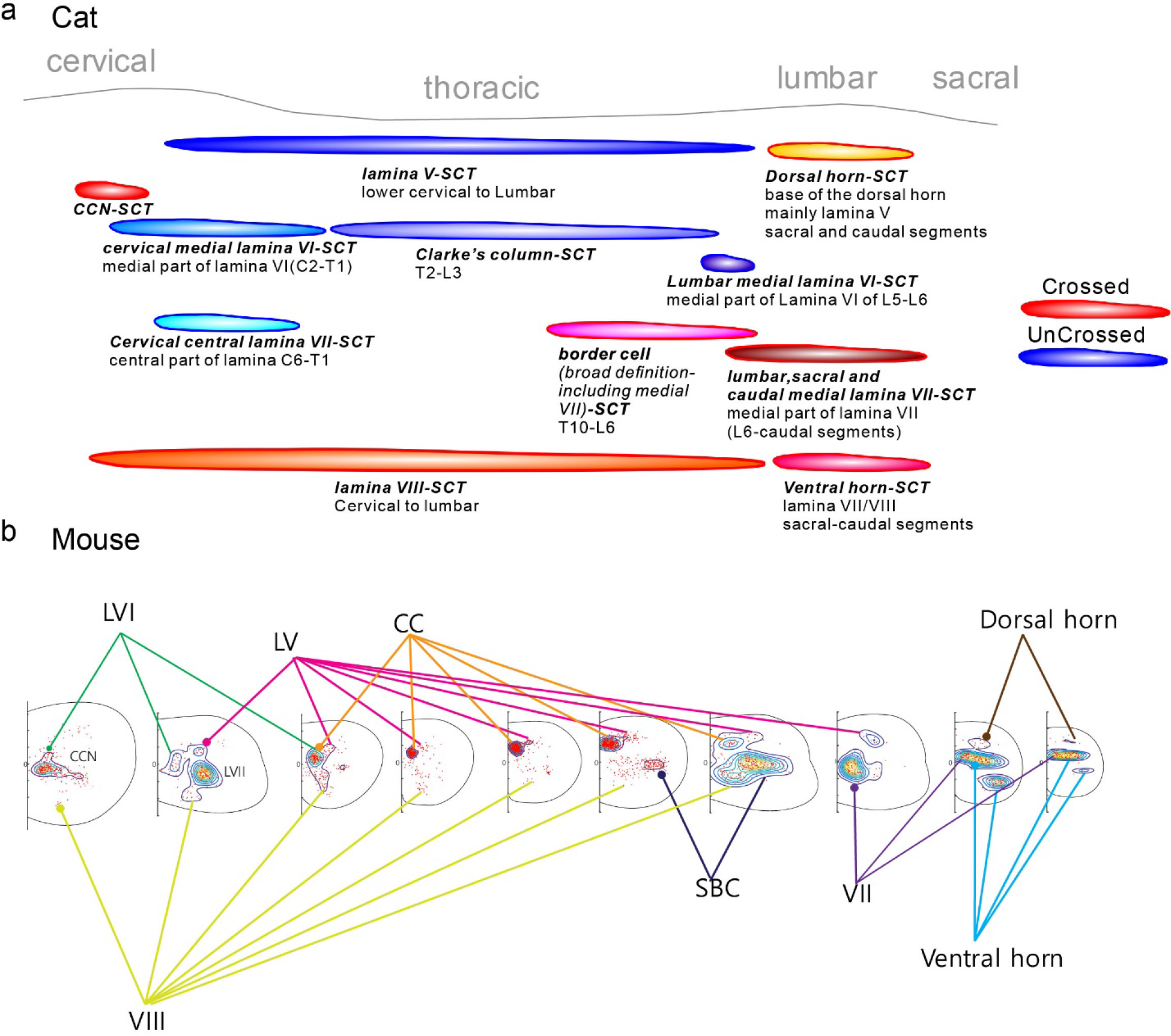
Diversity and anatomical location of SCTNs in cat and mouse. (a) Organization and diversity of SCTN subtypes described in cat. (b) Distribution of SCTN subtypes in early postnatal mice.

**Supplementary Fig. 2.**
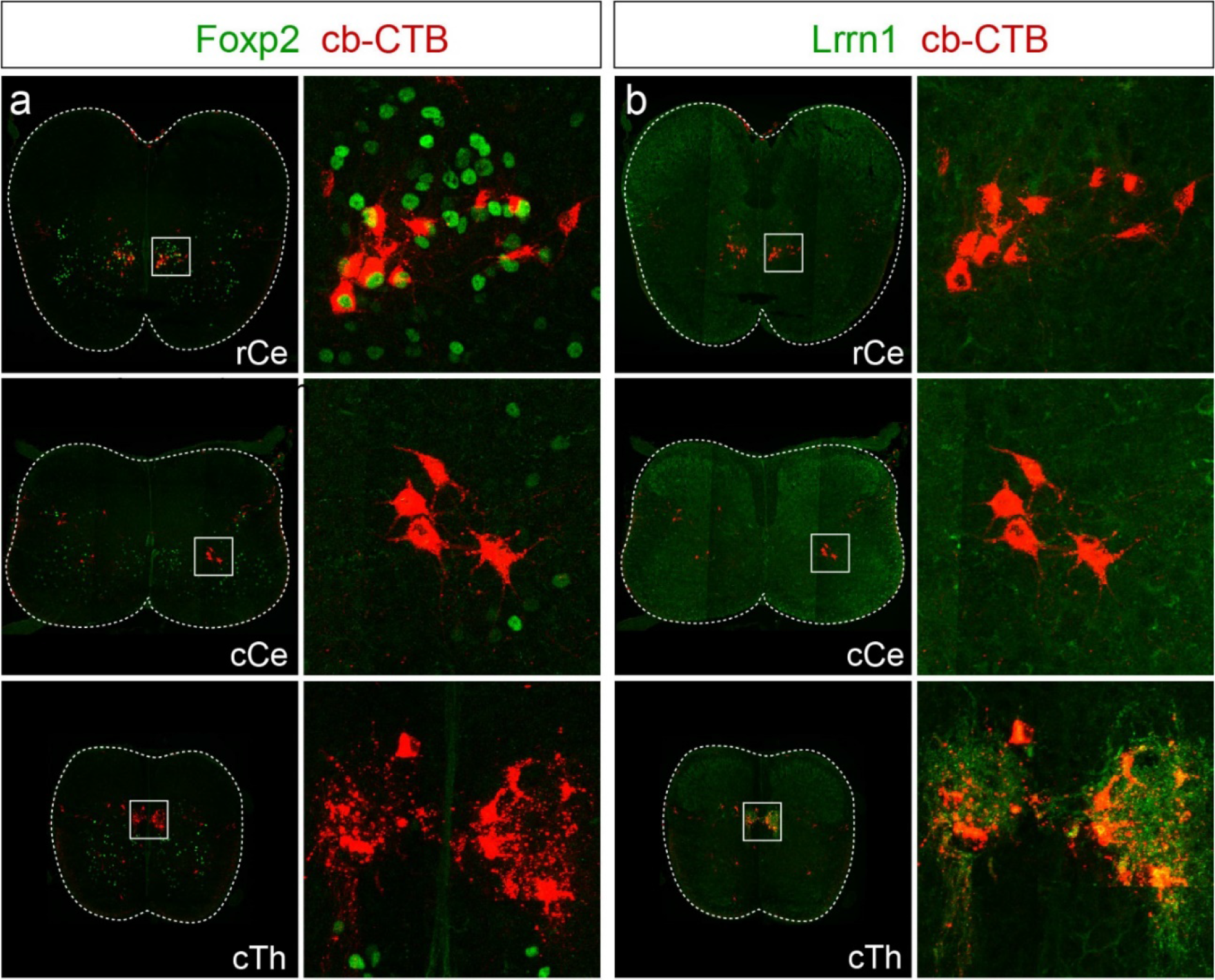
Expression of Foxp2 and Lrrn1 in SCTNs. (a) Expression of Foxp2 in retrogradely labeled SCTNs at rostral cervical (rCe), caudal cervical (cCe), and caudal thoracic (cTh) segments. Foxp2 is expressed in rCe SCTNs. (b) Expression of Lrrn1 in retrogradely labeled SCTNs at rCe, cCe, cTh segments. Lrrn1 is expressed in Th SCTNs.

**Supplementary Fig. 3.**
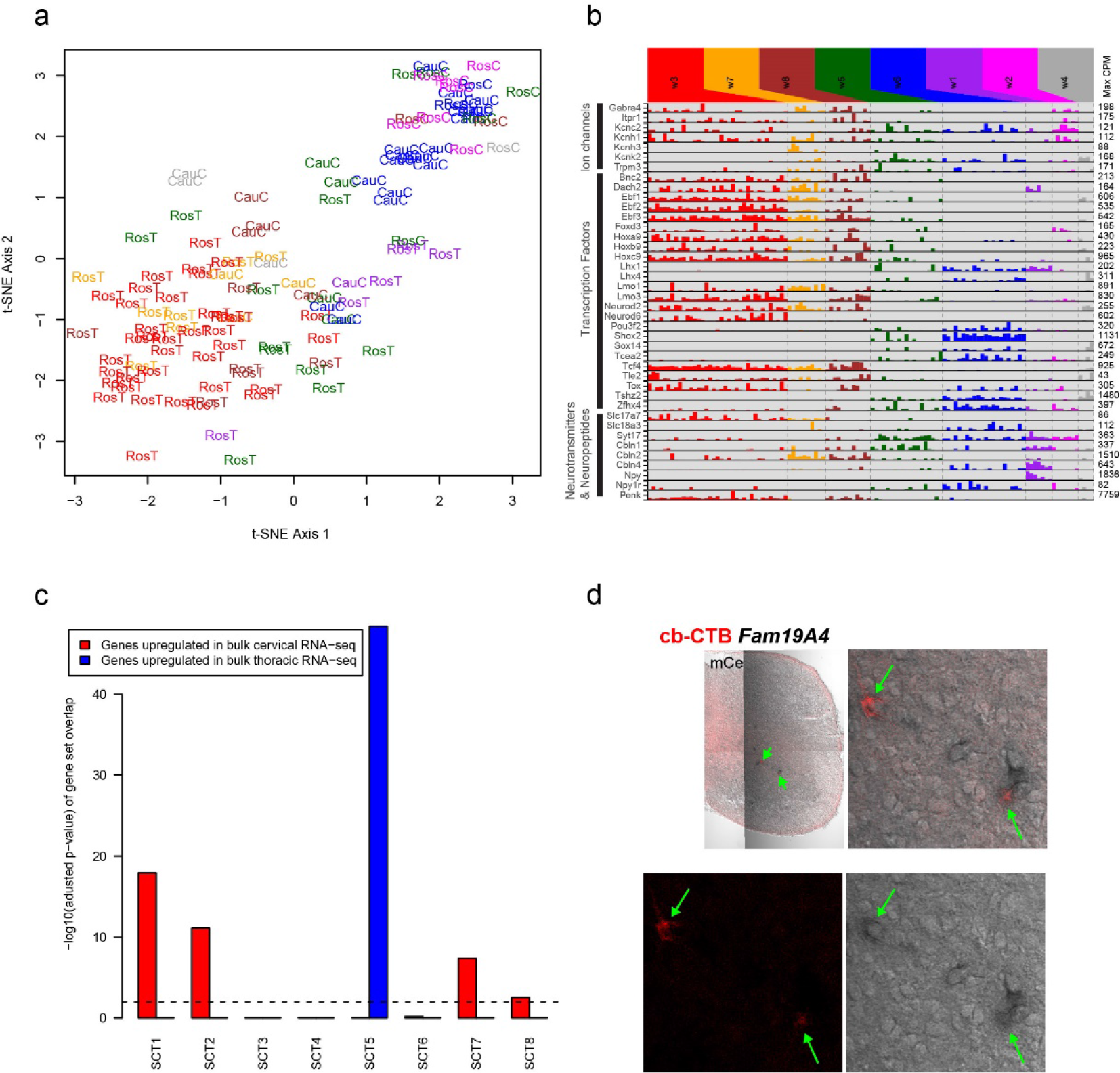
Analysis of SCTN scRNAseq data. (a) tSNE visualization of scRNAseq data from control cells, using only *Hox* genes for dimensionality reduction, showing segmental origin of single cells. Cells are colored by their original clusters (using all genes, as reflected in Fig. 3). RosT, rostral thoracic; RosC, rostral cervical; CauC, caudal cervical. (b) Barplot showing expression (TPM) values of selected ion channels, transcription factors, and neurotransmitters within clusters. (c) Comparison of bulk and scRNAseq data. The barplot shows the −log10 Bonferroni-adjusted p-value of the gene set overlap among cluster-specific genes from single-cell RNA-seq and differentially expressed genes from bulk RNA-seq of cervical and thoracic cells. Bulk RNA-seq genes were selected based on FDR<0.05 & fold-change>2. Cluster-specific gene sets from the single-cell RNA-seq were obtained for each cluster as follows: 1) for a given cluster, identify all genes upregulated in that cluster (FDR<0.05, fold-change>2) versus any other cluster using pairwise cluster comparisons, 2) select genes only if they are uniquely upregulated in that cluster i.e. no other cluster has significant upregulation of that gene with respect to any other cluster. P-values for each gene set overlap (8 cluster gene sets x 2 bulk gene = 16 overlaps in total) were calculated using the hypergeometric distribution, with the background gene set comprising all genes with any detection (>0) in either the single-cell or the bulk RNA-seq data. Gene set overlap p-values were adjusted post-hoc using the Bonferroni correction. As shown in the barplot, SCT1, SCT2, SCT7, and SCT8-specific gene sets show significant (p<0.01, dashed line) overlaps with genes upregulated in the bulk cervical RNA-seq data, whereas SCT5 expresses genes preferentially upregulated in the bulk thoracic RNA-seq data. (d) Validation of *Fam19A4* as cLVII SCTN marker. *Fam19A4* in situ hybridization (shown in black) was performed on Cb-CTB traced tissue sections (shown in red). Top left panel is a composite of tiled images.

**Supplementary Fig. 4.**
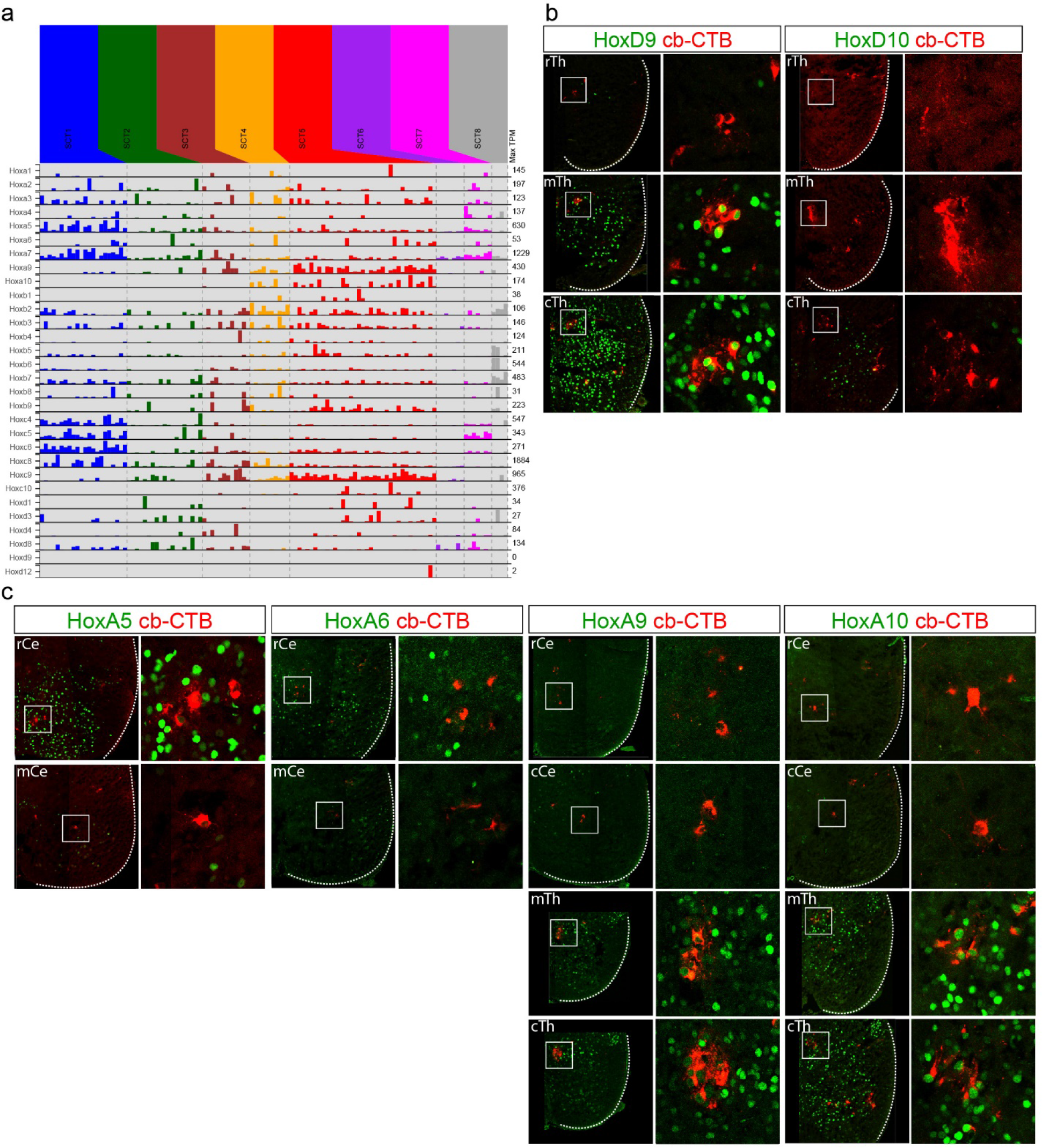
Hox expression in SCTNs. (a) Barplot showing the expression (TPM) of *Hox* genes from each of the four clusters in the control scRNAseq dataset, arranged by cluster identity (SCT1 through SCT8). (b) Expression of Hoxd9 and Hoxd10 in rTh, mTh, and cTh segments. SCTNs were labeled through CTB injection into the cerebellum. (c) Expression of indicated HoxA proteins in rCe, mCe, mTh, and cTh segments. SCTNs were labeled through CTB injection into the cerebellum.

**Supplementary Fig. 5.**
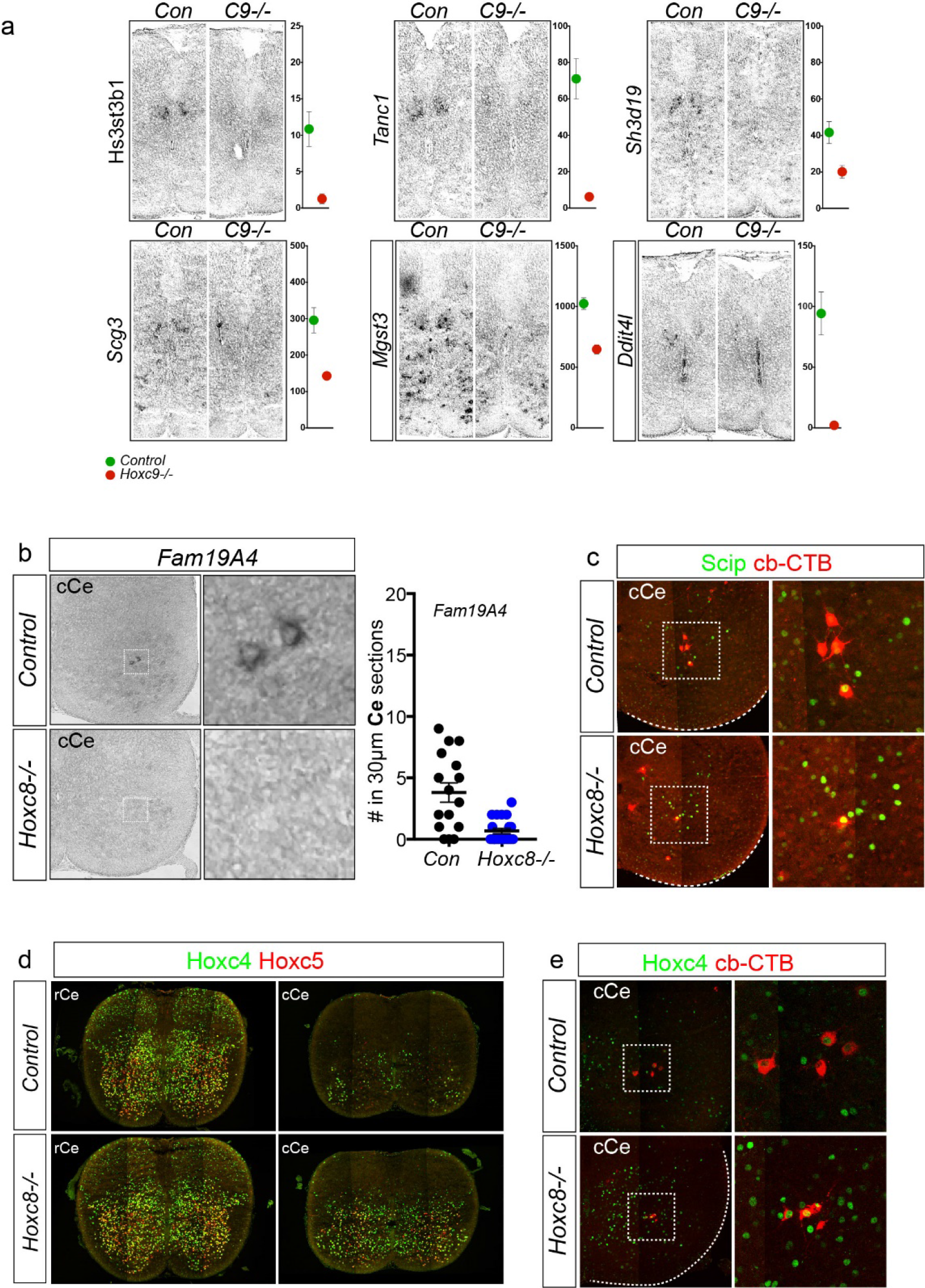
Analysis of SCNT specification in *Hoxc9* and *Hoxc8* mutants. (a) Analysis of CC neuron marker expression in thoracic segments of *Hoxc9* mutants. Expression levels from scRNAseq data in control and *Hoxc9−/−* thoracic SCTNs is shown on the right. (b) *Fam19A4* expression in *Hoxc8* mutants. Two tailed Student’s t test, p=0.0001 (***); Con, n=3(P0, 2; P6, 1; 16 sections); *Hoxc8−/−*, n=4(P0, 1; P6, 3; 21 sections). (c) Expression of Scip in SCTNs of *Hoxc8* mutants. (d) Hoxc4 and Hoxc5 protein expression at P0 in *Hoxc8* mutants. Both Hoxc4 and Hoxc5 are derepressed in cCe segments in *Hoxc8* mutants. (e) Caudal cervical SCTN ectopically express Hoxc4 in *Hoxc8* mutants.

**Supplementary Fig. 6.**
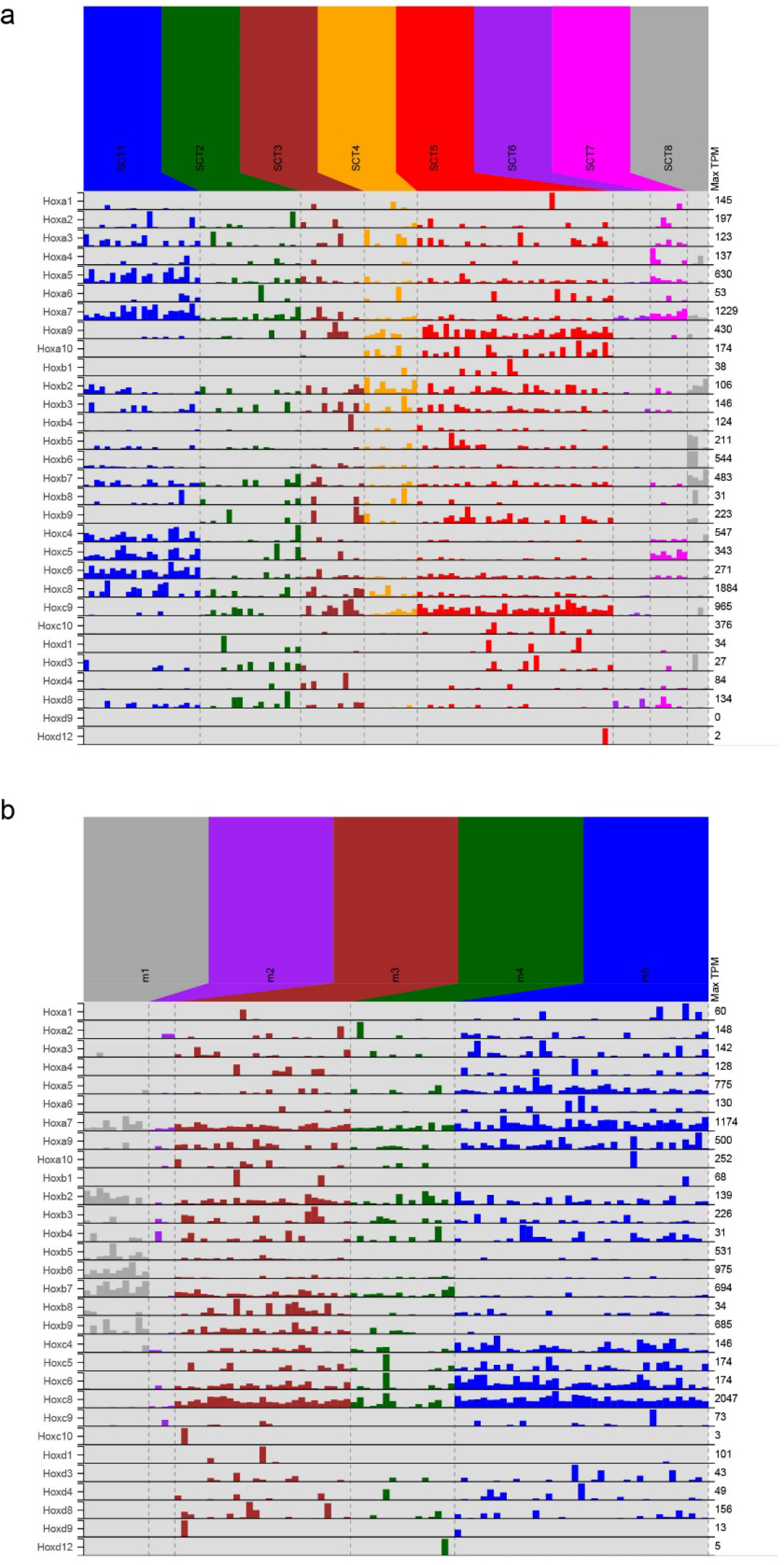
Expression of *Hox* genes in control and *Hoxc9* mutant SCTNs. (a) Barplot showing the expression (TPM) of *Hox* genes in the control scRNA-seq data, arranged by cluster identity (SCT1 through SCT8). Plot is identical to Supplementary Fig. 4a, and is shown here for direct comparison to *Hoxc9* mutants. (b) Same as panel A, but for *Hoxc9* mutant cells, arranged by mutant cluster identity. For both panels, *Hox* genes with no detected expression (in control or *Hoxc9* mutant cells) are not shown.

**Supplementary Fig. 7.**
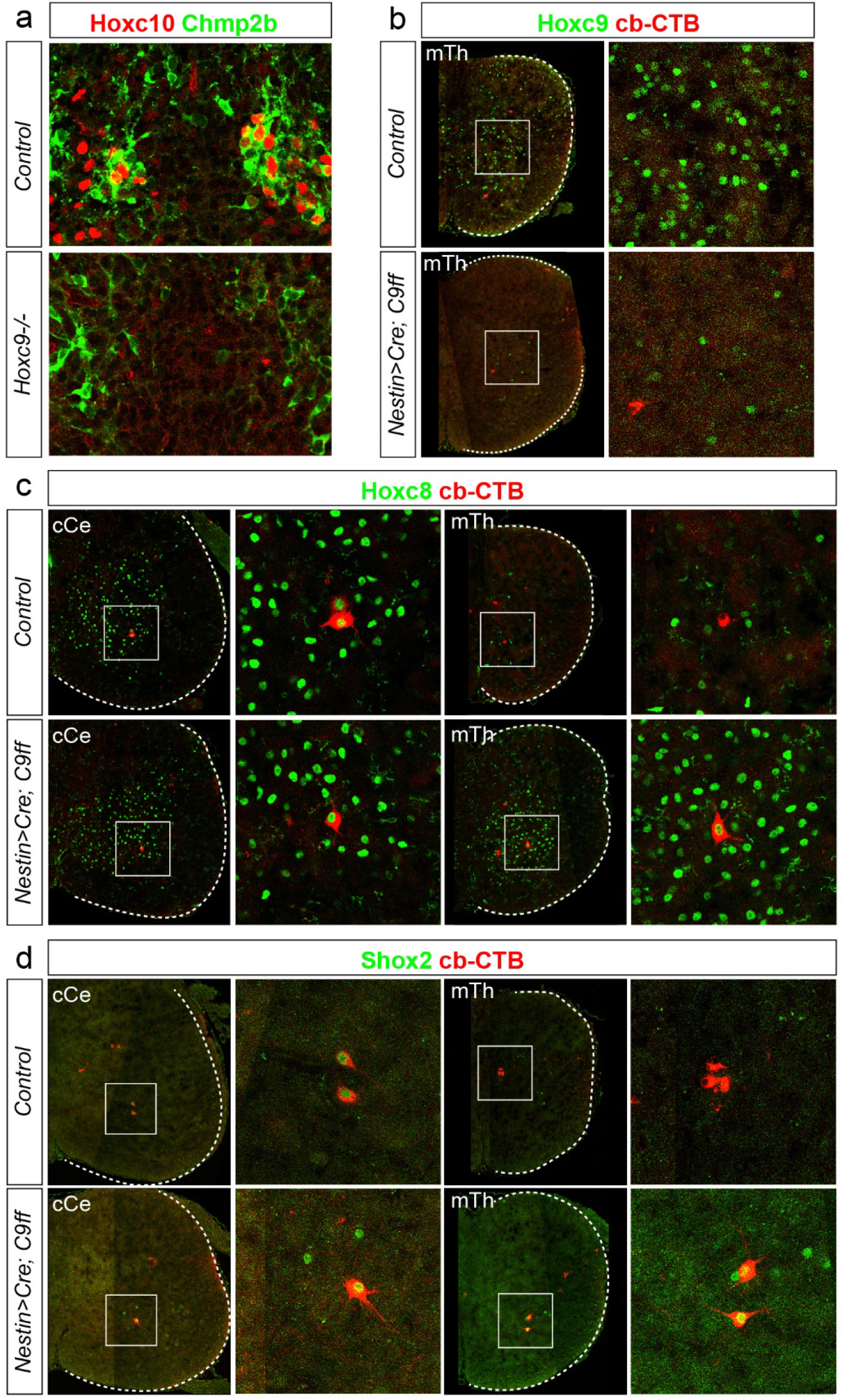
Analysis of SCTN differentiation in *Nestin::Cre; Hoxc9 flox/flox* mice. (a) Loss of Hoxc10 protein expression in thoracic segments of *Hoxc9* mutants. Chmp2b marks CC neurons. (b) Loss of Hoxc9 protein expression in thoracic segments of *Nestin::Cre; Hoxc9 flox/flox* mice. (c) In *Hoxc9* conditional mutants, SCTNs at rostral thoracic levels acquire Hoxc8 expression. (d) Rostral thoracic SCTNs ectopically express Shox2 in *Nestin::Cre; Hoxc9 flox/flox* mice.

**Supplementary Fig. 8.**
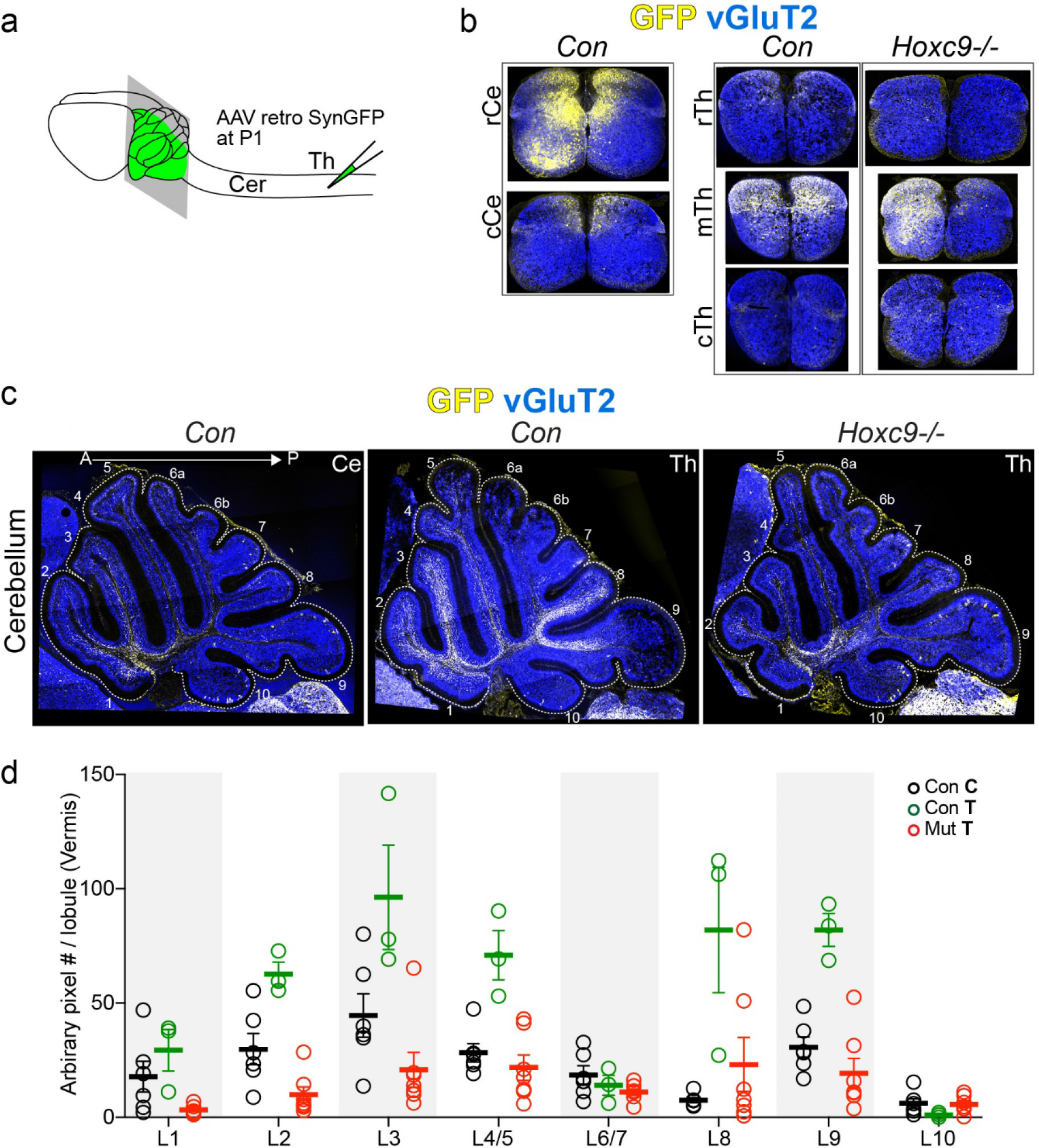
Analysis of spinal projections to cerebellum in *Hoxc9* mutants. (a) Strategy for spinal cord injection of *AAV-retro::SynGFP*. Virus was injected at P1 and GFP signals were examined at P6. (b) Analysis of GFP expression within indicated spinal cord sections of control and *Hoxc9* mutant mice. (c) Analysis of GFP expression in the cerebellum. Cerebellar lobule numbers are indicated. (d) Quantification of projections within the vermis. In Hoxc9 mutants there is a relatively marked reduction in the innervation of lobule 8. GFP pixel number were counted. Con Ce, n=5 mice; Con Th, 3 mice; *Hoxc9 −/−* (Mut) Th, n=5 mice.

## References

1 Bosco, G. & Poppele, R. E. Proprioception from a spinocerebellar perspective. Physiological reviews 81, 539–568, doi:10.1152/physrev.2001.81.2.539 (2001).

2 Tuthill, J. C. & Azim, E. Proprioception. Curr Biol 28, R194–R203 (2018).

3 Dietz, V. Proprioception and locomotor disorders. Nature reviews 3, 781–790 (2002).

4 Ghez, C., Gordon, J. & Ghilardi, M. F. Impairments of reaching movements in patients without proprioception. II. Effects of visual information on accuracy. Journal of neurophysiology 73, 361–372, doi:10.1152/jn.1995.73.1.361 (1995).

5 Gordon, J., Ghilardi, M. F. & Ghez, C. Impairments of reaching movements in patients without proprioception. I. Spatial errors. Journal of neurophysiology 73, 347–360, doi:10.1152/jn.1995.73.1.347 (1995).

6 Abelew, T. A., Miller, M. D., Cope, T. C. & Nichols, T. R. Local loss of proprioception results in disruption of interjoint coordination during locomotion in the cat. Journal of neurophysiology 84, 2709–2714, doi:10.1152/jn.2000.84.5.2709 (2000).

7 Akay, T., Tourtellotte, W. G., Arber, S. & Jessell, T. M. Degradation of mouse locomotor pattern in the absence of proprioceptive sensory feedback. Proceedings of the National Academy of Sciences of the United States of America 111, 16877–16882, doi:10.1073/pnas.1419045111 (2014).

8 Windhorst, U. Muscle proprioceptive feedback and spinal networks. Brain Res Bull 73, 155–202, doi:10.1016/j.brainresbull.2007.03.010 (2007).

9 Chen, H. H., Hippenmeyer, S., Arber, S. & Frank, E. Development of the monosynaptic stretch reflex circuit. Current opinion in neurobiology 13, 96–102 (2003).

10 Popova, L. B., Ragnarson, B., Orlovsky, G. N. & Grant, G. Responses of neurons in the central cervical nucleus of the rat to proprioceptive and vestibular inputs. Archives italiennes de biologie 133, 31–45 (1995).

11 Arsenio Nunes, M. L. & Sotelo, C. Development of the spinocerebellar system in the postnatal rat. The Journal of comparative neurology 237, 291–306, doi:10.1002/cne.902370302 (1985).

12 Matsushita, M. & Gao, X. Projections from the thoracic cord to the cerebellar nuclei in the rat, studied by anterograde axonal tracing. The Journal of comparative neurology 386, 409–421 (1997).

13 Matsushita, M., Hosoya, Y. & Ikeda, M. Anatomical organization of the spinocerebellar system in the cat, as studied by retrograde transport of horseradish peroxidase. The Journal of comparative neurology 184, 81–106, doi:10.1002/cne.901840106 (1979).

14 Sengul, G., Fu, Y., Yu, Y. & Paxinos, G. Spinal cord projections to the cerebellum in the mouse. Brain Struct Funct 220, 2997–3009, doi:10.1007/s00429-014-0840-7 (2015).

15 Edgley, S. A. & Grant, G. M. Inputs to spinocerebellar tract neurones located in stilling’s nucleus in the sacral segments of the rat spinal cord. The Journal of comparative neurology 305, 130–138, doi:10.1002/cne.903050112 (1991).

16 Kuno, M., Munoz-Martinez, E. J. & Randic, M. Sensory inputs to neurones in Clarke’s column from muscle, cutaneous and joint receptors. The Journal of physiology 228, 327–342 (1973).

17 Knox, C. K., Kubota, S. & Poppele, R. E. A determination of excitability changes in dorsal spinocerebellar tract neurons from spike-train analysis. Journal of neurophysiology 40, 626–646, doi:10.1152/jn.1977.40.3.626 (1977).

18 Osborn, C. E. & Poppele, R. E. The extent of polysynaptic responses in the dorsal spinocerebellar tract to stimulation of group I afferent fibers in gastrocnemius-soleus. J Neurosci 8, 316–319 (1988).

19 Hantman, A. W. & Jessell, T. M. Clarke’s column neurons as the focus of a corticospinal corollary circuit. Nature neuroscience 13, 1233–1239, doi:10.1038/nn.2637 (2010).

20 Philippidou, P. & Dasen, J. S. Hox genes: choreographers in neural development, architects of circuit organization. Neuron 80, 12–34, doi:10.1016/j.neuron.2013.09.020 (2013).

21 Sweeney, L. B. et al. Origin and Segmental Diversity of Spinal Inhibitory Interneurons. Neuron 97, 341–355 e343, doi:10.1016/j.neuron.2017.12.029 (2018).

22 Hayashi, M. et al. Graded Arrays of Spinal and Supraspinal V2a Interneuron Subtypes Underlie Forelimb and Hindlimb Motor Control. Neuron 97, 869–884 e865, doi:10.1016/j.neuron.2018.01.023 (2018).

23 Betley, J. N. et al. Stringent specificity in the construction of a GABAergic presynaptic inhibitory circuit. Cell 139, 161–174, doi:10.1016/j.cell.2009.08.027 (2009).

24 Shrestha, S. S. et al. Excitatory inputs to four types of spinocerebellar tract neurons in the cat and the rat thoraco-lumbar spinal cord. The Journal of physiology 590, 1737–1755, doi:10.1113/jphysiol.2011.226852 (2012).

25 Mann, M. D. Clarke’s column and the dorsal spinocerebellar tract: a review. Brain Behav Evol 7, 34–83, doi:10.1159/000124397 (1973).

26 Jung, H. et al. Global control of motor neuron topography mediated by the repressive actions of a single hox gene. Neuron 67, 781–796, doi:10.1016/j.neuron.2010.08.008 (2010).

27 Dasen, J. S., Liu, J. P. & Jessell, T. M. Motor neuron columnar fate imposed by sequential phases of Hox-c activity. Nature 425, 926–933, doi:10.1038/nature02051 (2003).

28 Bermingham, N. A. et al. Proprioceptor pathway development is dependent on Math1. Neuron 30, 411–422 (2001).

29 Rose, M. F., Ahmad, K. A., Thaller, C. & Zoghbi, H. Y. Excitatory neurons of the proprioceptive, interoceptive, and arousal hindbrain networks share a developmental requirement for Math1. Proceedings of the National Academy of Sciences of the United States of America 106, 22462–22467, doi:10.1073/pnas.0911579106 (2009).

30 Yuengert, R. et al. Origin of a Non-Clarke’s Column Division of the Dorsal Spinocerebellar Tract and the Role of Caudal Proprioceptive Neurons in Motor Function. Cell reports 13, 1258–1271, doi:10.1016/j.celrep.2015.09.064 (2015).

31 Bikoff, J. B. et al. Spinal Inhibitory Interneuron Diversity Delineates Variant Motor Microcircuits. Cell 165, 207–219, doi:10.1016/j.cell.2016.01.027 (2016).

32 Francius, C. et al. Identification of multiple subsets of ventral interneurons and differential distribution along the rostrocaudal axis of the developing spinal cord. PLoS One 8, e70325, doi:10.1371/journal.pone.0070325 (2013).

33 Dasen, J. S. Transcriptional networks in the early development of sensory-motor circuits. Current topics in developmental biology 87, 119–148, doi:10.1016/S0070-2153(09)01204-6 (2009).

34 Mendell, L. M. & Henneman, E. Terminals of single Ia fibers: distribution within a pool of 300 homonymous motor neurons. Science 160, 96–98 (1968).

35 Mendelsohn, A. I., Simon, C. M., Abbott, L. F., Mentis, G. Z. & Jessell, T. M. Activity Regulates the Incidence of Heteronymous Sensory-Motor Connections. Neuron 87, 111–123, doi:10.1016/j.neuron.2015.05.045 (2015).

36 Pecho-Vrieseling, E., Sigrist, M., Yoshida, Y., Jessell, T. M. & Arber, S. Specificity of sensory-motor connections encoded by Sema3e-Plxnd1 recognition. Nature 459, 842–846, doi:10.1038/nature08000 (2009).

37 Arber, S., Ladle, D. R., Lin, J. H., Frank, E. & Jessell, T. M. ETS gene Er81 controls the formation of functional connections between group Ia sensory afferents and motor neurons. Cell 101, 485–498 (2000).

38 de Nooij, J. C., Doobar, S. & Jessell, T. M. Etv1 inactivation reveals proprioceptor subclasses that reflect the level of NT3 expression in muscle targets. Neuron 77, 1055–1068, doi:10.1016/j.neuron.2013.01.015 (2013).

39 Inoue, K. et al. Runx3 controls the axonal projection of proprioceptive dorsal root ganglion neurons. Nature neuroscience 5, 946–954, doi:10.1038/nn925 (2002).

40 Surmeli, G., Akay, T., Ippolito, G. C., Tucker, P. W. & Jessell, T. M. Patterns of spinal sensory-motor connectivity prescribed by a dorsoventral positional template. Cell 147, 653–665, doi:10.1016/j.cell.2011.10.012 (2011).

41 Vrieseling, E. & Arber, S. Target-induced transcriptional control of dendritic patterning and connectivity in motor neurons by the ETS gene Pea3. Cell 127, 1439–1452, doi:10.1016/j.cell.2006.10.042 (2006).

42 Baek, M., Pivetta, C., Liu, J. P., Arber, S. & Dasen, J. S. Columnar-Intrinsic Cues Shape Premotor Input Specificity in Locomotor Circuits. Cell reports 21, 867–877, doi:10.1016/j.celrep.2017.10.004 (2017).

43 Bangma, G. C. & Donkelaar, H. J. T. Afferent Connections of the Cerebellum in Various Types of Reptiles. Journal of Comparative Neurology 207, 255–273, doi:DOI 10.1002/cne.902070306 (1982).

44 Gonzalez, A., ten Donkelaar, H. J. & de Boer-van Huizen, R. Cerebellar connections in Xenopus laevis. An HRP study. Anat Embryol (Berl) 169, 167–176 (1984).

45 Tourtellotte, W. G. & Milbrandt, J. Sensory ataxia and muscle spindle agenesis in mice lacking the transcription factor Egr3. Nature genetics 20, 87–91, doi:10.1038/1757 (1998).

46 Catela, C., Shin, M. M., Lee, D. H., Liu, J. P. & Dasen, J. S. Hox Proteins Coordinate Motor Neuron Differentiation and Connectivity Programs through Ret/Gfr alpha Genes. Cell reports 14, 1901–1915, doi:10.1016/j.celrep.2016.01.067 (2016).

47 Dasen, J. S., Tice, B. C., Brenner-Morton, S. & Jessell, T. M. A Hox regulatory network establishes motor neuron pool identity and target-muscle connectivity. Cell 123, 477–491, doi:10.1016/j.cell.2005.09.009 (2005).

48 Jung, H. et al. The Ancient Origins of Neural Substrates for Land Walking. Cell 172, 667–682 e615, doi:10.1016/j.cell.2018.01.013 (2018).

49 Hempel, C. M., Sugino, K. & Nelson, S. B. A manual method for the purification of fluorescently labeled neurons from the mammalian brain. Nat Protoc 2, 2924–2929, doi:10.1038/nprot.2007.416 (2007).

